# Med12 cooperates with multiple differentiation signals to facilitate efficient lineage transitions in embryonic stem cells

**DOI:** 10.1101/2024.01.22.576603

**Authors:** Max Fernkorn, Christian Schröter

## Abstract

Cell differentiation results from coordinated changes in gene transcription in response to combinations of signals. FGF, Wnt, and mTOR signals regulate the differentiation of pluripotent mammalian cells towards embryonic and extraembryonic lineages, but how these signals cooperate with general transcriptional regulators is not fully resolved. Here, we report a genome-wide CRISPR screen that reveals both signaling components and general transcriptional regulators for differentiation-associated gene expression in mESCs. Focusing on the Mediator subunit *Med12* as one of the strongest hits in the screen, we show that it regulates gene expression in parallel to FGF and mTOR signals. Loss of *Med12* is compatible with differentiation along both the embryonic epiblast and the extraembryonic primitive endoderm lineage, but impairs pluripotency gene expression and slows down transitions between pluripotency states. These findings suggest that *Med12* helps pluripotent cells to efficiently execute transcriptional changes during differentiation, thereby modulating the effects of a broad range of signals.

## Introduction

Cell differentiation during development is regulated by a suite of extracellular signaling systems that trigger expression changes of large gene modules. To execute such complex transcriptional responses, the activity of intracellular signaling effectors must be integrated with that of general transcriptional regulators that direct the activity of RNA polymerase II. How the signaling systems associated with pluripotency and differentiation work together with general transcriptional regulators during early lineage transitions is still not fully resolved.

The earliest cell differentiation events of mammalian embryogenesis first segregate the extraembryonic trophoblast from the inner cell mass (ICM). In a second step, the ICM further differentiates into extraembryonic primitive endoderm (PrE) and the pluripotent embryonic epiblast (Epi), which ultimately forms the fetus (Chazaud & Yamanaka, 2016). Subsequently, epiblast cells transition from the naïve pluripotent state at pre-implantation to formative and then primed pluripotency as they prepare for germ layer differentiation (Kalkan & Smith, 2014; Nichols & Smith, 2009; Smith, 2017). Embryonic stem cells (ESCs) allow modeling both the differentiation towards an extraembryonic PrE identity, as well as transitions between different pluripotent states (Kalkan et al., 2017). Mouse ESCs (mESCs) can either be maintained in medium containing serum and the cytokine LIF, or in a ground state of pluripotency using serum-free N2B27 medium supplemented with LIF and two small molecule inhibitors that activate Wnt/beta-Catenin signaling and inhibit FGF/ERK signaling, respectively (2i + LIF; Ying et al., 2008). Efficient PrE differentiation from mESCs can be achieved from ground state pluripotency by the forced expression of GATA transcription factors together with active FGF/ERK signaling (Fujikura et al., 2002; Schröter et al., 2015; Wamaitha et al., 2015). Transitions between pluripotency states in contrast can be triggered by the removal of small molecule inhibitors from the culture medium alone (Kalkan et al., 2017; Neagu et al., 2020). Together with experiments in the mouse embryo, these stem cell models have provided a comprehensive picture of the signaling control of the early lineage transitions in the mammalian embryo. Both PrE differentiation and exit of epiblast cells from naïve towards formative and primed pluripotency require FGF/ERK signaling as well as Wnt signaling inhibition (Athanasouli et al., 2023; Chazaud & Yamanaka, 2016; Kang et al., 2013; Nichols et al., 2009). PrE differentiation further benefits from LIF signaling (Morgani & Brickman, 2015), whereas the progression of epiblast cells is promoted by the Notch signaling effector RBPJ (Kalkan et al., 2019), and the mTOR signaling effector TFE3 (Betschinger et al., 2013; Villegas et al., 2019).

Developmental signaling systems often culminate in the activation or deactivation of sequence-specific transcription factors. In eukaryotes, transmission of such transcription factor activity changes into altered RNA polymerase activity at specific promoters requires large multiprotein assemblies such as the Mediator complex that physically bridges between transcription factors and the basal transcriptional machinery (Soutourina, 2018). The mammalian Mediator complex is formed by up to 30 subunits. It can be subdivided into a head, a core and a tail domain, and the transiently associated CDK8 module which consists of four subunits: the CDK8 kinase, CCNC, MED12, and MED13 (Luyties & Taatjes, 2022; Soutourina, 2018). Unperturbed Mediator function is required for expression of most protein-coding genes (Soutourina, 2018). Still, individual Mediator subunits have been linked to transcriptional changes in response to specific signaling systems. Deletion of *Sur2,* which encodes the MED23 subunit, for example abrogates transcriptional activation downstream of ERK-MAPK signaling in mESCs (Stevens et al., 2002). The CDK8 module in particular has been implicated in directing rapid changes in gene expression patterns in response to various stimuli, such as serum stimulation (Donner et al., 2010; Luyties & Taatjes, 2022). Furthermore, CDK8 inhibition in mouse and human ESCs impairs pluripotency exit, mirroring effects of MEK/ERK inhibition (Lynch et al., 2020). MED12, which activates CDK8 function in the kinase module (Knuesel et al., 2009; Park et al., 2018), is essential for axis elongation and the activation of Wnt target genes during mouse development (Rocha et al., 2010). Together, these works suggest that the transmission of specific developmental signals to RNA polymerase II activity can be mapped to specific Mediator subunits. How general these mappings are or whether they are context-dependent is however not clear. It is also not known how interference with Mediator activity in pluripotent cells affects differentiation efficiency of different lineages.

Here, we aim at identifying factors that mediate transcriptional changes in response to signaling events during early mammalian cell differentiation, using the expression of a *Spry4^H2B-Venus^* reporter allele as read-out in a genome-wide CRISPR screen. This reporter is an established indicator of developmental FGF/ERK signals, and its expression is switched on in both Epi and PrE cells during preimplantation development (Morgani et al., 2018). Our screen returns both known and new signaling inputs into *Spry4^H2B-Venus^* reporter expression, as well as several components of the Mediator and Elongator complexes. Using epistasis analysis, we demonstrate that *Med12*, one of the strongest hits in the screen, functions independently of and in parallel to the FGF and mTOR signaling systems in pluripotent cells. Functional assays showed that, while not strictly required for lineage transitions, loss of *Med12* leads to impaired signal responsiveness during pluripotency transitions. Collectively, these results point to new signal-independent functions of *Med12* that help cells to efficiently execute lineage transitions in early development.

## Results

### A genome-wide CRISPR screen identifies signaling and transcriptional regulators of *Spry4* expression

The differentiation of pluripotent cells towards different lineages rests on transcriptional changes triggered by extracellular signals. To identify effectors of signal-regulated gene expression during cellular differentiation, we performed a genome-wide CRISPR screen using the expression of a Sprouty4 reporter as a read-out (Hanna & Doench, 2020; Morgani et al., 2018; Fig. 1A). We chose Sprouty4 because it is a known target gene of developmental FGF signals, and shows strong expression changes upon both Epi and PrE differentiation (Morgani et al., 2018; Fig. 1 Supp 1). In serum + LIF medium, the *Spry4* reporter is expressed due to paracrine FGF signals. We generated Cas9-expressing *Spry4*:H2B-Venus reporter cells and transduced them with the Brie gRNA library targeting protein-coding genes (Doench et al., 2016). To identify positive and negative regulators of *Spry4* reporter expression, we flow sorted cells with decreased and increased fluorescence at 6 and 9 days after transduction, and determined enriched guides in sorted fractions (Fig. 1A). We first analyzed perturbations leading to reduced reporter expression. Gene-targeting gRNAs were more strongly enriched in the sorted fractions compared to non-targeting controls (Fig. 1B, Fig. 1 Supp 2, Supp Table 1). We used the robust rank algorithm (RRA; W. Li et al., 2014) to combine information from multiple gRNAs and to rank genes in each of the four conditions (1% and 5% gate at both day 6 and day 9, Supp Table 1). This analysis revealed up to 17 individual genes with an FDR ≤ 0.05, and up to 26 genes with an FDR ≤ 0.2 (Fig. 1C, Fig. 1 Supp 2). We compiled a list of hits by selecting genes that were detected with an FDR of ≤ 0.05 in one condition, or with an FDR of ≤ 0.2 in at least two conditions (Fig. 1D). Protein-protein interaction network analysis with String-DB revealed that most of our hits fell into a small number of groups that were highly connected and associated with specific molecular functions (Szklarczyk et al., 2015; Fig. 1E). One of these groups contained the FGF signaling genes *Fgfr1, Grb2, Sos1,* and *Ptpn11*, as would be expected given the strong regulation of *Spry4* by FGF signaling. Another group contained several genes involved in protein glycosylation and specifically the synthesis of heparan sulfates, such as *Slc35b2*, *Ext1*, *Ext2*, and *Extl3*. Heparan sulfates are crucial co-factors for efficient FGF signaling (Ornitz & Itoh, 2015). Since pooled CRISPR screens detect mainly cell-autonomous functions of gene perturbations, the appearance of these hits indicates that surface-tethered heparan sulfates determine a cell’s responsiveness to FGF signaling. Three further groups contained genes associated with ribosome biogenesis and translation (*Rpl9*, *Rpl18*, *Eif3i* and *Dhx37)*, as well as genes of the Elongator (*Elp2 – 6*, *Ikbkap* and *Kti12*) and Mediator complexes (*Med10, Med12, Med16, Med24* and *Med25*, Fig. 1E), raising the possibility that these factors could have gene- or signal-specific functions in mESCs.

**Fig. 1:**
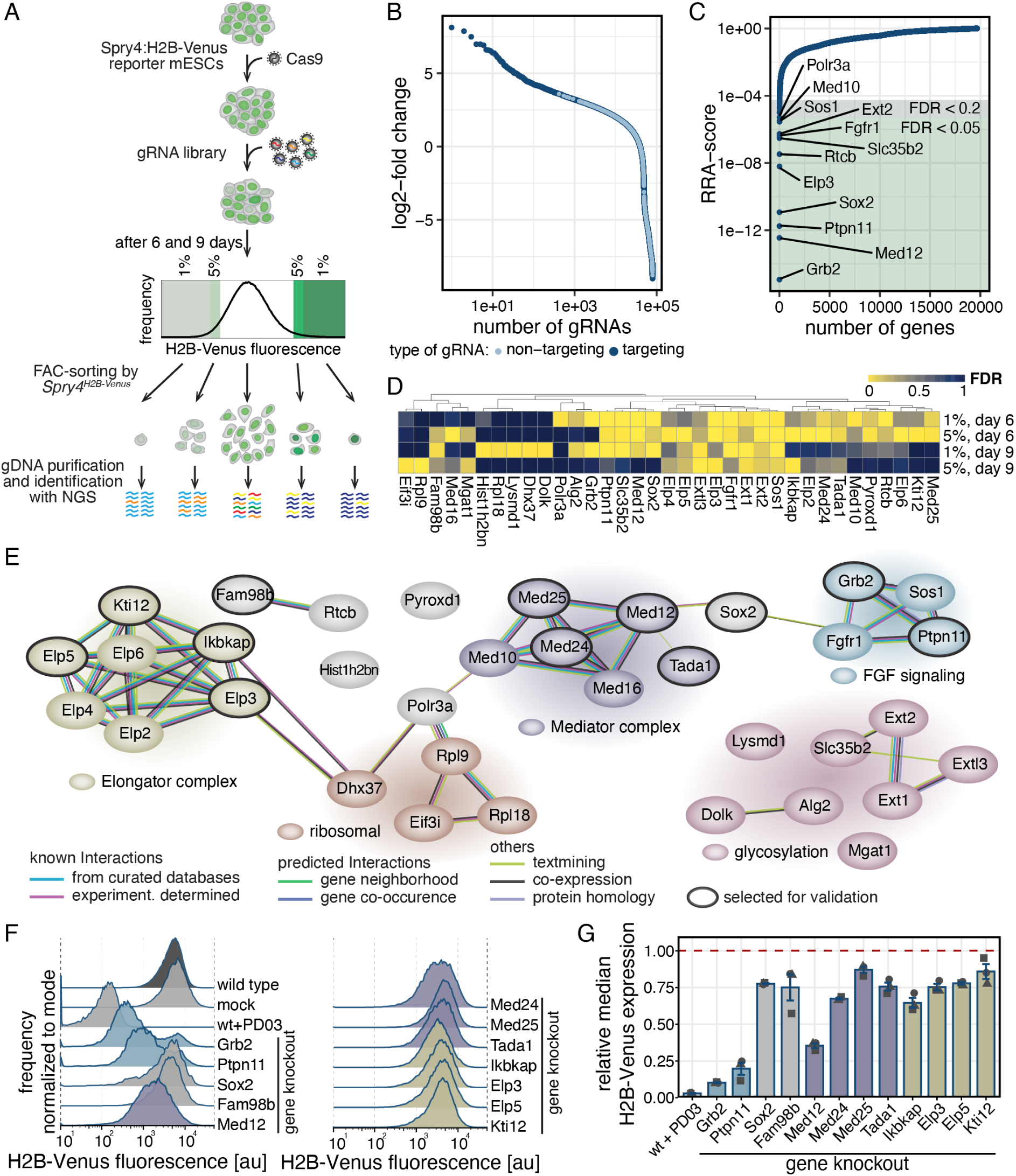
Genome-wide CRISPR knockout screen reveals positive regulators of Sprouty4 expression. **A** Schematic of the pooled CRISPR knockout screen. Cas9-expressing *Sprouty4^H2B-Venus/+^* reporter cells were transduced with a genome-wide gRNA library targeting protein-coding genes (Doench et al., 2016). Cells with in- or decreased fluorescence were flow sorted 6 or 9 days after transduction, and gRNAs enriched in sorted populations identified by sequencing. **B**, **C** Enrichment of gene-targeting (dark blue) and control gRNAs (light blue) in cells sorted for the lowermost 1% of H2B-Venus signal 6 days after gRNA transduction, displayed as log2-fold change (B) or RRA score of corresponding genes (C). Green and gray background in (C) indicates FDR < 0.05 and < 0.2, respectively. **D** Hierarchical clustering of gene perturbations leading to reduced *Spry4^H2B-Venus^* expression (FDR ≤ 0.05 in at least one condition or FDR ≤ 0.2 in at least two conditions). **E** Protein interaction network of genes shown in (D) based on String-DB. Background colors of genes and gene clusters were manually assigned based on classification by functional similarity. **F**, **G** H2B-Venus expression in *Spry4^H2B-Venus/+^* reporter cells upon knockout of selected candidate genes 6 d after transfection. (F) shows histograms of one representative experiment, (G) shows mean ± SEM of median H2B-Venus expression from N = 3 independent experiments, normalized to fluorescence levels in control wild-type cells (dashed red line). p < 0.05 for wild type vs. *Elp3*, *Elp5*, *Fam98b*, *Ikbkap*, *Kti12*, *Med25* or *Tada1* knockouts; p < 0.01 for wild type vs. *Grb2*, *Med12*, *Med24*, *Ptpn11*, *Sox2* knockouts or PD03-treated cells (Benjamin-Hochberg-adjusted, one-sided, paired t-test).

Next, to validate selected hits in an independent experimental setting, and to evaluate their effect size, we knocked out individual candidate genes using the most enriched gRNA from the screen. Since we were interested in mechanisms of transcriptional regulation, we focused on hits related to the Mediator and Elongator complexes, but also included *Grb2* and *Ptpn11* as a reference to evaluate the effects of perturbing FGF signal transduction, and *Sox2* and *Fam98b* as candidates that could not clearly be linked to a functional group (Fig. 1E). Flow cytometry showed that knockout of all tested candidates led to a reduction of mean *Spry4*:H2B-Venus fluorescence levels, albeit to a different degree: Knockout of components of the Elongator complex as well as *Fam98b* and *Sox2* affected Sprouty4:H2B-Venus expression only mildly, whereas knockout of the FGF signaling genes *Grb2* and *Ptpn11* had the strongest effect, although they did not reach the reduction achieved via pharmacological inhibition of FGF signal transduction with the MEK inhibitor PD0325901 (PD03) (Fig. 1F, G). Knockout of Mediator components reduced *Spry4*:H2B-Venus levels to different degrees, with *Med12* having the strongest effect, reducing *Spry4*:H2B-Venus levels to 35.3% ± 1.6% compared to control (Fig. 1F, G). Thus, our screen and validation establish *Med12* as a strong candidate regulator of signal-dependent gene expression in mESCs.

We next sought to use our screen to identify negative regulators of *Spry4* expression, by looking at perturbations that led to increased reporter expression (Fig. 1A, right side of histogram). Also here, gene-targeting gRNAs were enriched over control guides in the sorted fractions (Fig. 2 Supp 1A-D, Supp Table 1). Ordering according to RRA-scores revealed up to 12 and 24 genes with FDR values of ≤ 0.05 and ≤ 0.2, respectively in each of the conditions (Fig. 2A, Fig. 2 Supp 1E-G, Supp Table 1). Using the same criteria as for the positive regulators, we compiled a list of 29 potential negative regulators of *Spry4* transcription (Fig. 2B). Analysis with String-DB again showed that many of these hits were highly connected and associated with specific molecular functions (Fig. 2C). A large group of genes encoded proteins that localized to mitochondria, or were otherwise associated with mitochondrial functions. Several genes had signaling functions: *Lztr1* is a negative regulator of RAS-MAPK and hence FGF signaling, in line with the strong representation of genes promoting FGF signaling amongst the positive regulators of *Spry4* expression. We detected four genes that were related to mTOR signaling (*Tsc1*, *Tsc2*, *Flcn*, and *Lamtor*). Finally, we found a big group that contained genes involved in chromatin modification and transcription, amongst them three genes encoding SWI/SNF related proteins. Thus, similarly to the collection of positive hits identified above, our screen yields both signaling and transcriptional regulators that negatively control *Spry4* expression. To select hits for validation, we focused on genes associated with mTOR signaling, because these genes have also been implicated in the maintenance of pluripotency (Betschinger et al., 2013; M. Li et al., 2018; Villegas et al., 2019). We also included *Lztr1* as an FGF signaling gene, and *Smarcc1* as a representative of the group of chromatin modifiers. Knockout of all six individual candidate genes with single gRNAs led to an increase of mean *Spry4*:H2B-Venus fluorescence levels in reporter cells. The effect of knocking out the mTOR signaling genes was stronger than that of knocking out *Lztr1* or *Smarcc1*, and almost doubled reporter expression levels compared to mock-transfected cells (Fig. 2D, E).

**Fig. 2:**
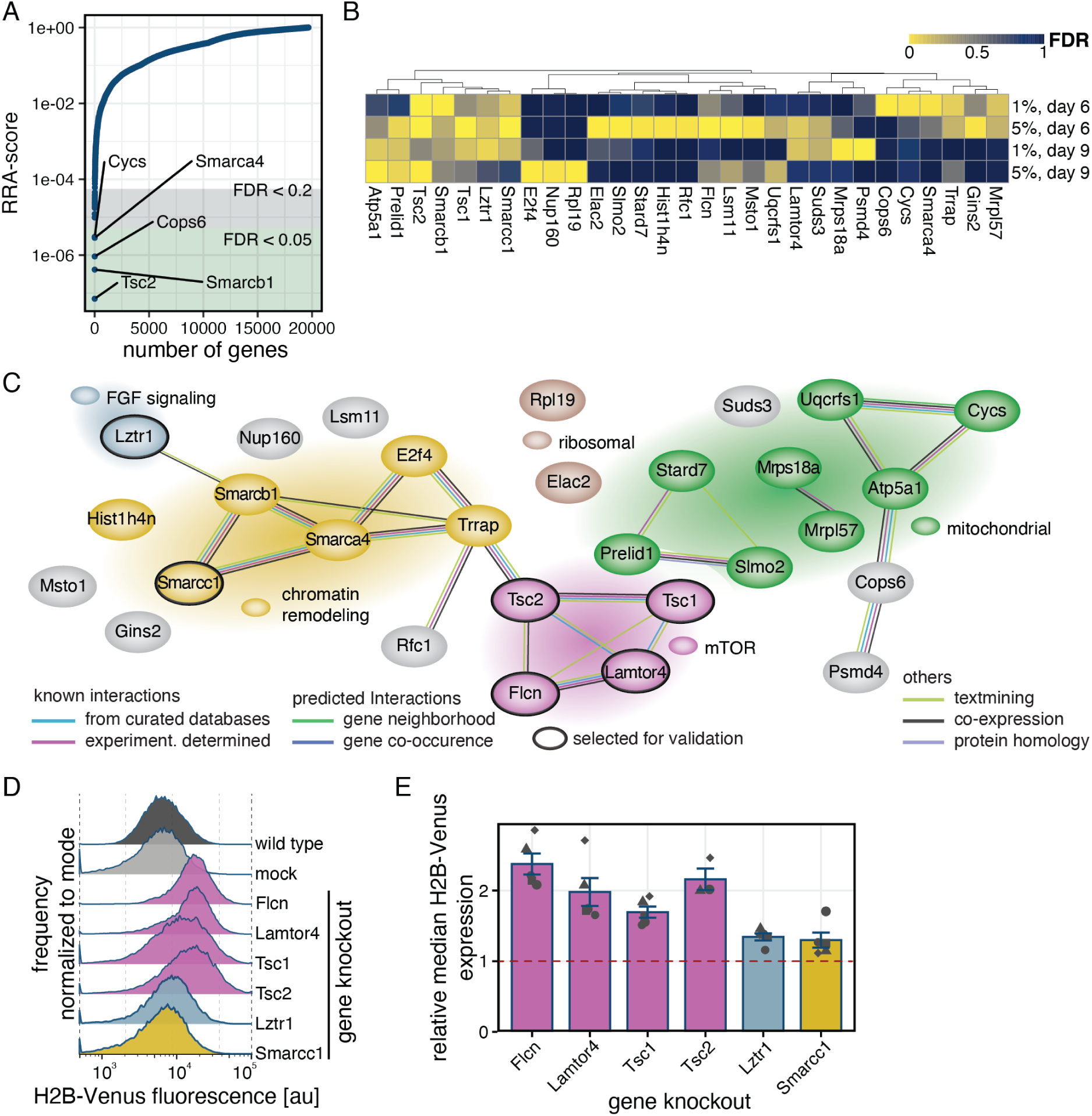
Genome-wide CRISPR knockout screen reveals negative regulators of Sprouty4 expression. **A** RRA scores of genes corresponding to gRNAs enriched in cells sorted for the topmost 1% of H2B-Venus signal on day 6 after gRNA transduction. **B** Hierarchical clustering of gene perturbations leading to increased *Spry4^H2B-Venus^* expression (FDR ≤ 0.05 in at least one condition or FDR ≤ 0.2 in at least two conditions). **C** Protein interaction network of genes shown in (B) based on String-DB. Background colors of genes and gene clusters were manually assigned based on classification by functional similarity. **D**, **E** H2B-Venus expression in *Spry^H2B-Venus/+^* reporter cells upon knockout of selected candidate genes 6 d after transfection. (D) shows histograms of one representative experiment, (E) shows mean ± SEM of median H2B-Venus expression, N ≥ 3 independent experiments, normalized to fluorescence levels in control cells transfected with a non-targeting control (dashed red line). p < 0.05 for mock-transfected wild type vs. *Smarcc1* knockout; p < 0.01 for mock-transfected wild type vs. all other knockouts (Benjamin-Hochberg-adjusted, one-sided, paired t-test).

### *Med12* regulates gene expression in mESCs independently from pluripotency-related signaling pathways

We then wanted to know how the signaling and transcriptional regulators identified in the CRISPR-screen were functionally related. We focused on *Med12*, since of all transcriptional regulators, it had the strongest effect on *Spry4* reporter expression. Furthermore, MED12 is a component of Mediator’s kinase module which could couple the activities of specific signaling systems to transcriptional activity (Lynch et al., 2020; Rocha et al., 2010). To analyze *Med12* functions in mESCs, we first generated monoclonal *Med12*-mutant cell lines. We generated multiple independent clonal lines lacking part of exon 7, a region that was also targeted by sgRNAs used in the CRISPR-screen, and confirmed the loss of MED12 protein expression by immunoblotting (Fig. 3 Supp 1A, B). *Med12*-mutant cells grew normally in both serum + LIF and 2i + LIF medium, and showed a reduced increase in *Spry4* reporter expression upon switching from 2i + LIF to the N2B27 base medium (Fig. 3 Supp 1C, D). ppERK levels were indistinguishable between wild-type and *Med12*-mutant lines (Fig. 3 Supp 1E, F), indicating that reduced *Spry4* reporter expression is not due to deregulated FGF/ERK signaling. Fluorescent in-situ hybridization revealed lower numbers of *Spry4* mRNAs in *Med12*-mutant compared to wild-type cells Fig. 3 Supp 1G, H), suggesting that *Med12* regulates reporter expression transcriptionally. Intriguingly, we also found slightly lower numbers of *Nanog* mRNAs in *Med12* mutants in 2i medium, indicating that *Med12* could be quantitatively involved pluripotency gene expression.

**Fig. 3:**
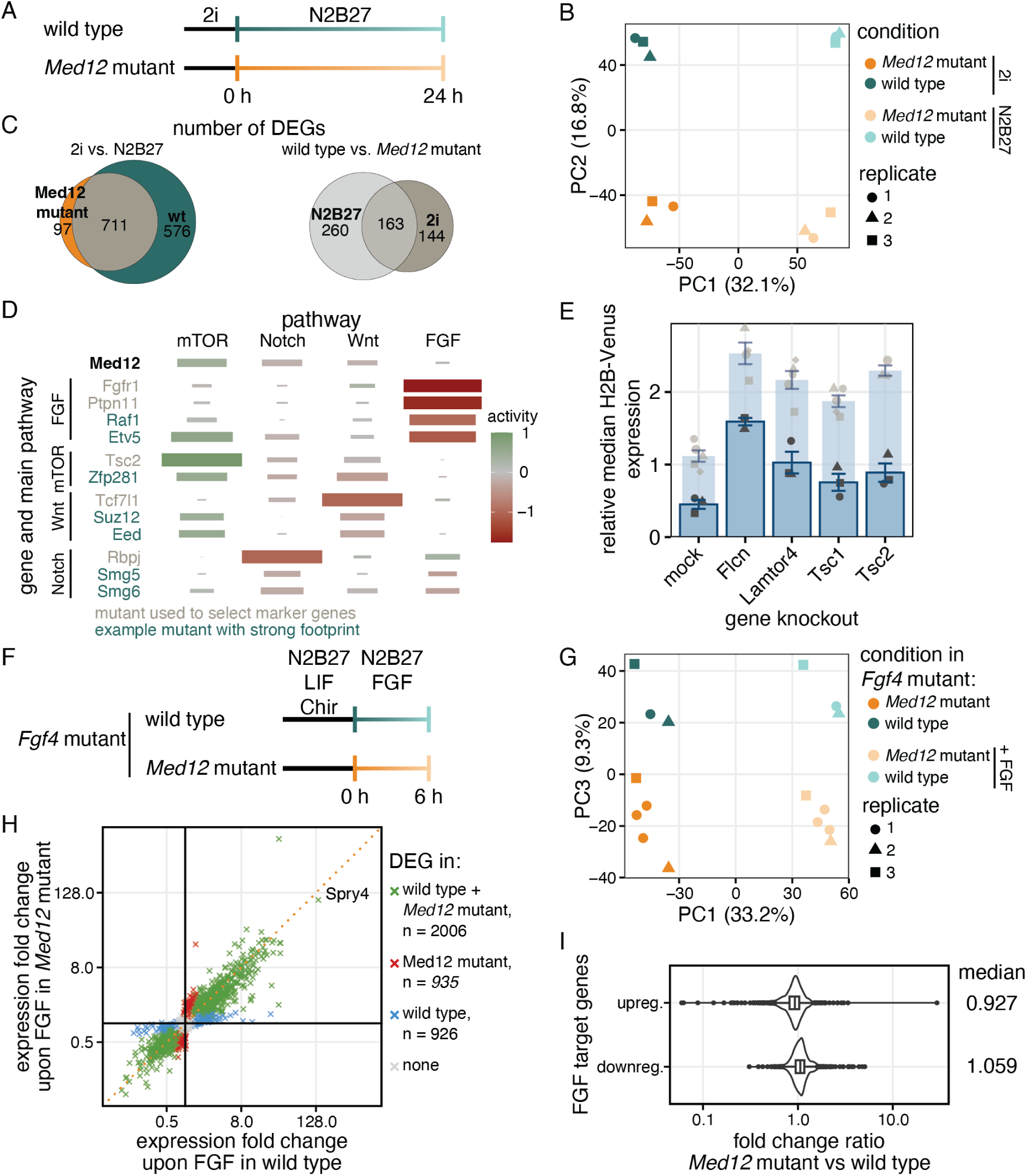
*Med12* affects gene expression independently of pluripotency related signaling systems. **A** Schematic of experiment to identify *Med12*-regulated genes by bulk RNA sequencing. **B** Principal component analysis transcriptomes from (A). **C** Euler-diagram showing the number differentially expressed genes (log2-fold change > |1|, adjusted p-value < 0.01) in bulk transcriptomes. Left panel compares genes differentially expressed upon 24 h of differentiation between *Med12-*mutant and wild-type cells, right panel compares genes differentially expressed upon loss of *Med12* between N2B27 and 2i. **D** Expression footprint analysis using a set of 50 marker genes per pathway defined in (Lackner et al., 2021). Top row shows footprint of *Med12*-mutant cells. Lower rows show expression footprints of mutants from Lackner et al., 2021, for comparison. Gray: Mutants that were used to select marker genes. Green: Independent example mutants that show a strong and specific footprint of one of the pathways. Tile color indicates relative pathway activity, tile size indicates spearman correlation of the expression of footprint genes with pathway-defining mutants (Lackner et al., 2021). **E** Median H2B-Venus fluorescence upon mutation of mTOR related genes in *Med12*-mutant *Spry4^H2B-Venus/+^* cells, normalized to H2B-Venus expression in *Med12* wild-type cells. Median H2B-Venus fluorescence upon mutation of mTOR related genes in *Med12* wild-type cells is reproduced from Fig. 2E for comparison (light blue). Error-bars indicate SEM, points individual replicates, N ≥ 3 independent experiments. **F** Schematic of experiment to test *Med12*-dependency of FGF target genes by RNA sequencing. **G** Principal component analysis transcriptomes from (F). PC2 (not shown; 12.8% of variance) separated experimental replicates from each other. **H** Gene expression fold changes and number of significantly differentially expressed genes (adjusted p-value < 0.01) upon FGF4 stimulation in wild-type versus *Med12*-mutant cells. Dotted orange line indicates the unity line. **I** Ratio of expression fold changes for FGF target genes between wild-type and *Med12*-mutant cells, for up- (top) and downregulated genes (bottom). FGF target genes were defined as having a log2-fold change in wild-type cells upon FGF4 stimulation > |1| and an adjusted p-value < 0.05.

To assess gene expression differences between wild-type and *Med12-*mutant cells more globally, we therefore performed bulk RNA sequencing of cells in 2i medium and upon 24 h of differentiation in N2B27 (Fig. 3A). Principal component analysis separated samples in pluripotency and differentiation medium along PC1 (32.1% of variance), and wild-type and *Med12-*mutant cells along PC2 (16.8% of variance, Fig. 3B). Fewer genes were differentially expressed between pluripotency conditions and 24 h in N2B27 in *Med12*-mutant compared to wild-type cells. Most genes that were differentially expressed in the mutant were also differentially expressed in the wild type (Fig. 3C, left, Supp Table 2). When comparing between wild-type and *Med12*-mutant cells within each culture condition, we found that more genes were differentially expressed between the two genotypes in N2B27 than in 2i (Fig. 3C, right, Supp Table 2). Thus, loss of *Med12* impairs gene expression changes at the exit from pluripotency. *Med12* has a closely related paralogue named *Med12l*. *Med12l* expression was upregulated in *Med12-*mutant cells (Fig. 3 Supp 2A), and *Spry4* reporter expression in *Med12-* mutant lines could be further reduced by simultaneously knocking-out *Med12l* (Fig. 3 Supp 2B). Thus, it is possible that *Med12l* partially compensates for loss of *Med12* function in the mutant lines.

Exit from pluripotency is regulated by the interplay of a set of signaling systems, such as mTOR, Notch, Wnt, and FGF (Betschinger et al., 2013; Kalkan et al., 2019; Ying et al., 2008). We next asked if gene expression downstream of any of these signaling systems was affected by loss of *Med12*, using previously defined sets of target genes specific for each signaling system (Lackner et al., 2021). A strong expression change of such a set of target genes between wild type and mutant cells can be considered a footprint of the perturbation of the corresponding signaling system. We quantified this footprint of the loss of MED12 by comparing the gene expression changes of the defined list of target genes during 24h of differentiation between the wild-type and the mutant. This gene expression footprint was strongest for mTOR target genes, less pronounced for Notch and Wnt target genes, and virtually absent for FGF target genes (Fig. 3D, top row). In comparison to previously described examples (Fig. 3D, lower rows, Lackner et al., 2021)) for signal-specific perturbations, the effects of the loss-of MED12 appeared less specific for a single pathway. We functionally tested the relationship between *Med12* and mTOR signaling by knocking out the mTOR signaling genes *Flcn*, *Lamtor4*, *Tsc1* and *Tsc2* in *Med12*-mutant *Spry4^H2B-Venus/+^* cells. Similarly to the situation in the wild type, knockout of these genes led to increased reporter expression, albeit from a lower baseline level (Fig. 3E). Therefore, *Med12* regulates *Spry4* expression independently from mTOR signaling. In addition to mTOR signaling genes and Mediator subunits, our CRISPR screen revealed a large number of genes involved in FGF signal transduction, but the footprinting analysis suggested that FGF inputs into gene expression were independent from *Med12*. To further corroborate this result, we generated *Med12* mutations in the background of an *Fgf4-*mutant mESC line, which allowed us to specifically analyze the effects of FGF signaling upon the switch from pluripotency to differentiation medium. We wanted to focus on immediate gene expression changes triggered by FGF signaling, and therefore analyzed gene expression changes by bulk RNA-sequencing after 6 h of transfer into N2B27 medium with or without addition of exogenous FGF (Fig. 3F). Again, principal component analysis showed that cells in pluripotency and differentiation medium were separated along PC1 (33.2% of variance), and wild-type and *Med12-*mutant cells separated along PC3 (9.3% of variance), both when analyzing independent *Med12*-mutant clonal cell lines as well as experimental replicates (Fig. 3G). If expression of FGF target genes was generally dependent on *Med12,* we would expect that their fold-change is lower in *Med12-* mutant compared to wild-type cells, and that the number of differentially expressed genes would be higher in the wild-type than in the *Med12*-mutant. However, when plotting the fold expression-change for each gene upon 6 h of FGF stimulation for wild-type versus *Med12*-mutant cells, we found that the majority of genes were induced to a similar degree in both genotypes (2006 genes), while similar numbers of only 926 and 935 genes were differentially expressed in the wild-type or the *Med12*-mutant only (Fig. 3H, Supp Table 3). Furthermore, the ratios of fold-change values of FGF target genes between *Med12*-mutant and wild-type cells showed a unimodal distribution with a mode of 1.068 for downregulated and 0.927 for upregulated genes (Fig. 3J). While this slight deviation of the mode from 1 leaves open the possibility that *Med12* influences their expression magnitude, overall these results argue against a strong and specific role of *Med12* in the regulation of FGF target genes.

### Extraembryonic endoderm differentiation in *Med12* mutant cells

Loss of the Mediator subunit Med24 compromises PrE differentiation (Hamilton et al., 2019). We therefore asked if *Med12* was likewise required for the segregation of Epi and PrE identities, using their differentiation of upon transient expression of GATA6-mCherry as a model system (Fig. 4A, Schröter et al., 2015). A mix of SOX17-positive PrE cells and NANOG-positive Epi cells differentiated irrespective of *Med12* status, but the fraction of PrE cells was lower, and that of Epi cells was higher in mutant compared to wild-type cultures (Fig. 4B, C). Those differences in cell type ratios could not be rescued by supplementing the differentiation medium with FGF4 (Fig. 4 Supp. 1A, B). However, similarly high proportions of PrE cells differentiated from both wild-type and *Med12* mutant cells when cells sorted for high GATA6-mCherry levels were treated with FGF4 during differentiation (Fig. 4 Supp. 1C, D), indicating that the main cause for these ratio differences were lowered expression levels of the GATA6-mCherry transgenes in *Med12* mutant cell lines compared to wild type (Fig. 4D). This conclusion was further corroborated when we connected GATA6-mCherry expression levels with differentiation outcome in individual cells through live-cell imaging followed by immunostaining for fate markers (Schröter et al., 2015). GATA6-mCherry levels rose more slowly and reached lower peak levels in the mutant relative to wild type, even when using longer induction times (Fig. 4 Supp 1F). Still, PrE differentiation could be predicted based on GATA6-mCherry expression levels with similar precision in both wild-type and *Med12* mutant cells (Fig. 4 Supp. 1 G – I), and GATA6-mCherry threshold values required to trigger PrE differentiation appeared even slightly lowered in *Med12* mutant cells (Fig. 4 Supp. 1 J). Thus, loss of *Med12* impairs transgene induction levels but not general PrE differentiation potential. To test if loss of *Med12* had more subtle effects on PrE differentiation, we performed single-cell RNA sequencing (scRNAseq) of wild-type and *Med12-*mutant cells in pluripotency conditions, and after GATA6-mCherry induction for 4 h (wild type) or 8 h (*Med12* mutants) followed by 20 h of differentiation (Fig. 4E). Cells from 2i + LIF (pluripotency) and N2B27 medium (differentiation) formed two distinct groups in UMAP space that were marked by expression of the Wnt/β-Catenin target Sp5 (pluripotency) or differentiation marker Dnmt3l, respectively (differentiation) (Fig. 4F, G). Within both the pluripotency and the differentiation group, wild type and *Med12*-mutant cells segregated from each other, indicating that loss of *Med12* affected single-cell transcriptional state in both conditions (Fig. 4F). In the differentiation group, the PrE markers Sox17, Dab2, and Cubn and the Epi markers Nanog and Fgf4 showed a mutually exclusive expression pattern in both wild-type and *Med12-*mutant cells (Fig. 4G), providing an opportunity to compare lineage-specific effects of the loss of *Med12* between Epi and PrE cells. To consistently identify Epi and PrE cells across genotypes, we integrated the five differentiated samples and clustered them to separate two major groups that were separated along in principle component space (Fig. 4H). *Gata6* and *Nanog* expression identified these two clusters as PrE (cluster 0) and Epi (cluster 1), respectively (Fig. 4H, I, Supp Table 4). The proportion of PrE cells in this dataset was again lower in *Med12* mutants compared to wild-type, consistent with immunostaining results (Fig. 4J). To evaluate the roles of *Med12* on lineage-specific gene expression changes upon differentiation, we first selected genes that had a log2 fold change ≥ 0.5 between pluripotency and differentiation in wild-type cells and plotted their fold-change in the mutant (Fig. 4K). These fold-changes showed a unimodal distribution in all four conditions (up- and downregulation, Epi and PrE differentiation). Furthermore, when plotting the fold-change of all genes upon differentiation against each other for wild-type and mutant cells, we found that data points uniformly clustered around the line of unity (Fig. 4 Supp 3 A, B). Finally, many of the top differentially expressed genes between wild type and mutant were the same in all in pluripotent, Epi, and PrE cells (Fig. 4 Supp 3 A, B, Supp Table 5). Taken together, these results suggest that loss of *Med12* does not affect specific gene modules during differentiation. but rather results in gene expression changes that are shared between differentiated states.

**Fig. 4:**
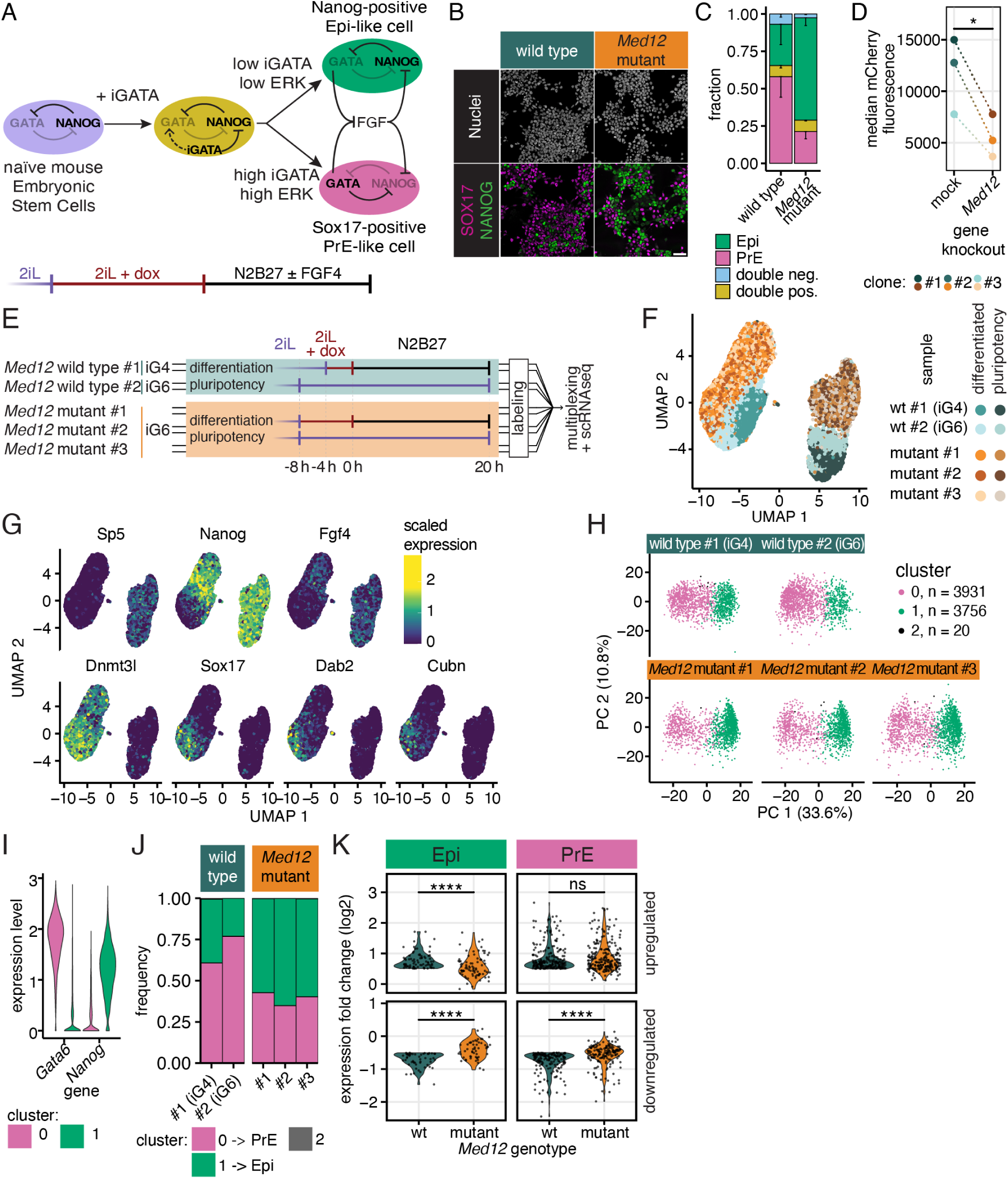
Role of *Med12* in PrE differentiation. **A** Schematic of experimental approach to model differentiation of mESCs towards epiblast and primitive endoderm via GATA induction. **B** Immunostaining for the Epi-marker NANOG (green) and the PrE marker SOX17 (magenta) after 8 h of GATA6 induction followed by 20 h of differentiation in wild-type and *Med12*-mutant cells. Scale bar: 50 µm. **C** Cell type proportions in wild-type and *Med12*-mutant cells differentiated as in (B). N = 3, n > 1100 cells per replicate, error bars indicate SEM. **D** Median Gata6-mCherry fluorescence upon 8h dox induction in three independent clonal GATA6-mCherry inducible cell lines 7 days after transfection with control or *Med12-*targeting gRNAs. * indicates p ≤ 0.05, paired student’s t-test. **E** Schematic of the single cell RNA sequencing experiment to compare single cell transcriptional signatures between wild-type and *Med12*-mutant cells in pluripotency and upon PrE differentiation. Two *Med12* wild-type lines carrying inducible GATA4- or GATA6-mCherry transgenes (iG4 and iG6), and three independent *Med12*-mutant cell lines derived from the GATA6-mCherry inducible line were included as replicates. Samples were multiplexed and pooled before mRNA capture and library preparation to minimize batch effects. Doxycycline induction was 4 h and 8 h in wild-type and *Med12*-mutant cells, respectively. **F** UMAP plot of single cell transcriptomes from all 10 samples. **G** Ln-scaled expression levels of selected marker genes projected onto the UMAP plot from (F). **H** Principal component analysis and Louvain-clustering of single-cell transcriptomes from wild-type and *Med12*-mutant cells after differentiation. **I** Ln-transformed expression levels of the PrE-marker gene *Gata6* and the Epi-marker gene *Nanog* in the differentiated samples split by cluster. Cluster 2 was excluded due to the small number of cells. **J** Proportions of cell types in the sequencing experiment, identified by clustering, in wild-type and *Med12*-mutant cells. **K** Comparison of up- (top panels) and downregulated genes (bottom panels) in wild-type and *Med12-* mutant cells upon differentiation from pluripotency to Epi (left) or PrE (right). Shown are genes with a log2-change of expression > 0.5 in wild-type cells. ns indicates p ≥ 0.05, **** indicates p ≤ 0.0001, paired Wilcoxon signed rank test. For clarity, measurements from the two *Med12* wild-type and the three *Med12*-mutant lines were pooled in panels I and K.

Prompted by the observation that GATA6-mCherry induction levels were consistently lowered in *Med12* mutant cells, we asked if loss of *Med12* had more global effects on transcriptional output and mRNA levels in cells. The multiplexed nature of our dataset allowed us to compare mRNA levels between conditions, revealing that differentiation led to an increase in captured mRNAs in both wild-type and mutant cells (Fig. 4 Supp. 3A). Furthermore, less mRNAs were captured from differentiated *Med12* compared to wild type cells (median number of UMIs 28227 and 25649, respectively, Fig. 4 Supp 3 A). While EU-labelling of newly synthesized RNAs suggested that higher mRNA content in differentiated compared to pluripotent cells were associated with increased transcriptional output, we could not detect significant differences in EU-labelling between wild-type and *Med12* mutant cells (Fig. 4 Supp 3 B – E). This leaves open the possibility that reduced mRNA levels in *Med12* mutant cells arise from effects other than reduced transcriptional output.

### *Med12* regulates dynamics of pluripotency transitions

Besides bridging between signals, sequence-specific transcription factors and the core transcriptional machinery, the Mediator complex also regulates enhancer-promoter interactions in ESCs (Kagey et al., 2010). We reasoned that impaired reconfiguration of these contacts might impact differentiation dynamics, and therefore tested how pluripotency transitions were affected by loss of *Med12.* We first probed exit from naïve pluripotency with a colony formation assay (Kalkan et al., 2017), where we differentiated cells in N2B27 for 48 h, followed by seeding at clonal density in 2i + LIF medium (Fig. 5A). Because we grew cells in the presence of LIF before transfer into N2B27, wild-type cells still formed a large number of pluripotent colonies (Lackner et al., 2021). In *Med12*-mutant cells in contrast, the number of pluripotent colonies was significantly reduced (Fig. 5B, left, Fig. 5 Supp 1A, Supp Table 6). To test how stable this phenotype is under different signaling conditions, we carried out the same colony assay in an *Fgf4* mutant background. Upon supplementation of N2B27 with FGF4, the number of pluripotent colonies formed by *Fgf4; Med12* double mutant cells was much reduced, both compared to the *Fgf4* single mutant supplemented with FGF4, and the *Fgf4; Med12* double mutant in the absence of FGF4 (Fig. 5B, right, Fig. 5 Supp 1B). Although we cannot rule out the possibility that the use of FBS required for the proliferation of *Fgf4* mutant cells in 2i + LIF medium (Raina et al., 2021) had selective effects on the *Med12* mutant cells, these results suggest that loss of *Med12* impairs the ability of single cells to re-establish a gene expression profile compatible with growth in 2i + LIF following transient differentiation in N2B27.

**Fig. 5:**
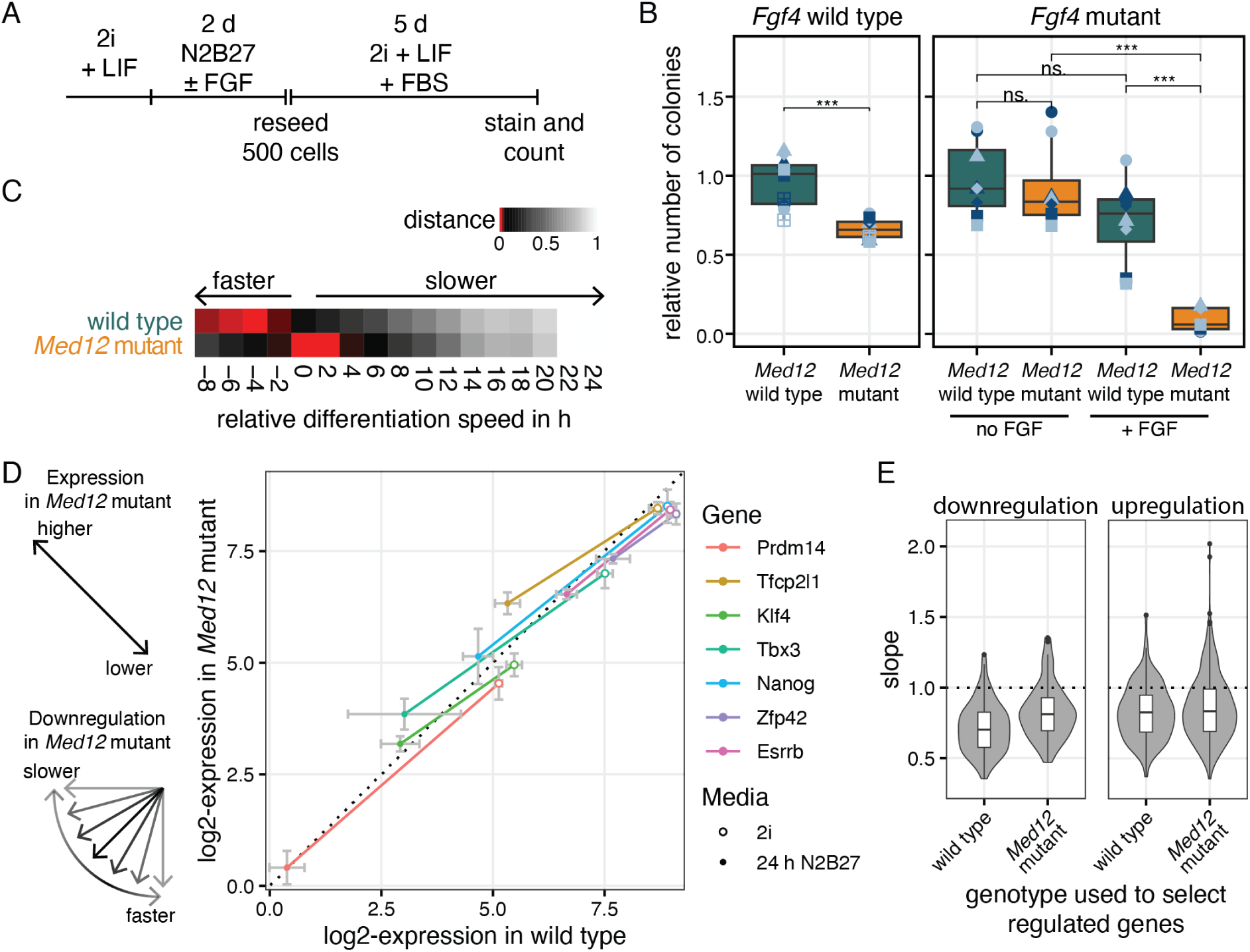
Mutation of *Med12* affects dynamics of pluripotency transitions. **A** Experimental approach to determine clonogenicity of *Med12*-mutant and wild-type cells. 2i + LIF was supplemented with FBS to support growth of *Fgf4*-mutant cells after reseeding. **B** Number of colonies after treatment as indicated in (A) for both wild-type and *Med12*-mutant cells in an *Fgf4* wild-type (left) and *Fgf4* mutant background with and without supplementation with 10 ng/µL FGF4 (right). N ≥ 4 independent experiments, same symbols in different colors indicate technical replicates within an independent experiment. ns indicates p ≥ 0.05, *** indicates p ≤ 0.001, paired Wilcoxon signed rank test. **C** Estimation of differentiation delay in wild-type and *Med12*-mutant cells, relative to a published time resolved gene expression dataset (Lackner et al., 2021). Plot shows the normalized Euclidean distance of the expression of naïve marker gene panel (*Prdm14*, *Tfcp2l1*, *Klf4*, *Tbx3*, *Nanog*, *Zfp42*, *Esrrb*) to the reference dataset. The negative delay values for our wild-type cells likely reflect small differences in experimental design compared to the study by (Lackner et al., 2021). **D** Expression of naïve pluripotency marker genes in 2i (open circles) and after 24 h differentiation in N2B27 (dots) in wild-type versus *Med12*-mutant cells. Relative expression values shown as log2(TPM), error bars indicate standard deviation. **E** Distribution of slopes of regulation (gene expression change over differentiation in wild type divided by gene expression change over differentiation in *Med12* mutant) for the 100 genes with the strongest negative (left panel) and positive (right panel) fold-change in wild-type and *Med12*-mutant cells. Bar indicates median, boxes indicate 25^th^ and 75^th^ percentile.

The ability to form pluripotent colonies in the colony formation assay depends on a cell’s pluripotency state at the beginning of the differentiation period, and its ability to react to changing culture conditions. We used our bulk RNA sequencing data comparing wild-type and *Med12*-mutant cells after 24 h in N2B27 to evaluate which of these properties were affected upon loss of *Med12*. We first determined differentiation delays of the two genotypes using a published high time-resolution reference dataset (Lackner et al., 2021). Surprisingly, this analysis indicated that *Med12*-mutant cells differentiated more slowly than the wild-type (Fig. 5C). This could be a consequence of stronger expression of naïve genes in pluripotency conditions, or reflect slower downregulation of these genes in N2B27 in the *Med12* mutant. To distinguish between these possibilities, we plotted the expression levels of 7 selected pluripotency marker genes in 2i and after 24 h in N2B27 in wild-type versus *Med12*-mutant cells. In this plot, data points above and below the unit line indicate higher and lower expression in the mutant relative to the wild type, respectively, and the slope of the connecting line between expression in 2i and N2B27 indicates the dynamics of down-regulation (Fig. 5D, left). We found that in 2i medium, all marker genes were less strongly expressed in the mutant compared to the wild type. Consistent with our results from bulk RNA sequencing, expression levels of the pluripotency genes in the single-cell RNA sequencing dataset (Fig. 4E) *Klf4*, *Zfp42*, and *Tbx3* were reduced in the mutant cells, whereas levels of *Prdm14*, *Nanog*, *Esrrb*, and *Tcf21l1* were similar between the two genotypes (Fig. 5 Supp 1C). Furthermore, their slopes during differentiation were consistently smaller than one (Fig. 5D). Such a systematic reduction in the slope of downregulation was also observed when we analyzed the top 100 downregulated genes in wild-type or *Med12*-mutant cells (Fig. 5E, left). Upregulation of genes is similarly affected, as the top 100 upregulated genes also showed a slope lower than 1 (Fig. 5E, right). Since a loss of Oct4 (POU5F1) can be sufficient to block the differentiation potential of mESCs (Radzisheuskaya et al., 2013), we checked the protein expression level by immunostaining (Figure 5 Supp 1C). However, there was no significant difference in expression levels between *Med12*-mutant and wild-type cells (Figure 5 Supp 1D). Taken together, these results indicate that *Med12* mutant cells show an impaired pluripotency gene expression profile as well as a slowed reconfiguration of gene expression programs upon changing culture conditions, thereby implicating *Med12* as a global regulator of pluripotency transition dynamics.

## Discussion

Here we use a *Spry4^H2B-Venus^* reporter to screen for regulators of developmental gene expression in pluripotent stem cells. This screen returned components of the FGF/ERK and the mTOR signaling systems that positively and negatively regulate reporter expression, respectively, as well as several members of the Mediator and Elongator complexes. Focusing on *Med12* we show that it cooperates with multiple signaling systems to regulate gene expression in pluripotent cells. *Med12* was not required for the differentiation of Epi and PrE identities, but dynamics of pluripotency transitions were impaired in *Med12-*mutant cells. Together, these results suggest that *Med12* amplifies transcriptional changes in pluripotent cells, possibly by contributing to Mediator’s function in establishing promoter-enhancer contacts (Kagey et al., 2010).

Previous genome-wide screens have used retention of clonogenicity or the continued expression of pluripotency-associated reporter alleles in differentiation conditions as read-outs to identify regulators of pluripotency and lineage transitions (Betschinger et al., 2013; Kagey et al., 2010; M. Li et al., 2018; Villegas et al., 2019). In our study, the combination of a *Spry4^H2B-Venus^* allele with flow sorting constitutes a highly sensitive read-out that is focused on the activity of specific signaling systems in pluripotent cells, and reliably detects new regulators even if they have only small effect sizes. The screen’s specificity shows in the strong representation of genes involved in the FGF/ERK signaling cascade, from genes that encode synthetases for FGF co-factors, over FGF receptors, to intracellular signaling genes. Surprisingly however, we did not find any sequence-specific transcription factors downstream of FGF/ERK signaling in our screen. This could be explained by the expression of multiple functionally redundant FGF/ERK signaling effectors in pluripotent cells, or through previously proposed transcription factor-independent regulation of RNA polymerase activity by ERK (Tee et al., 2014).

In addition to components of FGF and mTOR signaling systems, the screen returned several members of the SWI/SNF and the Mediator complexes. The core Mediator complex is thought to be required for the expression of most genes in eukaryotic genomes, but individual subunits have been suggested to regulate gene expression downstream of specific signaling systems such as the serum response network (Donner et al., 2010; Stevens et al., 2002), or Wnt (Rocha et al., 2010). However, when we tested this idea for *Med12*, we found that its loss did not phenocopy the effects of specific signaling perturbations. This finding suggests that previously reported functional connections, such as the link between *Med12* and Wnt signaling (Rocha et al., 2010), are strongly context-dependent.

*Med12* is a critical component of the Mediator-associated CDK8-module. In contrast to pharmacological inhibition of CDK8 which has been reported to boost pluripotency similarly to ERK signaling inhibition (Lynch et al., 2020), we find that loss of *Med12* leads to lower expression levels of pluripotency markers. These opposing phenotypes indicate that MED12 has functions that are independent from the CDK8-module (Aranda-Orgilles et al., 2016). This would also explain why we did not detect any other CDK8-module components in our screen. Lynch et al. found that maintenance of pluripotency required the presence of CDK8, but absence of its kinase activity. Another possible explanation for the opposing phenotypes of CDK8 inhibition and *Med12* loss-of-function therefore is that MED12 participates in assembling kinase-inactive CDK8 complexes that support pluripotency.

*Med12* has been found to maintain pluripotency not only in the epiblast, but also in the trophoectoderm (Halstead et al., 2024). Its role to maintain pluripotency gene expression reported here is supported by earlier studies which found that MED12 and NANOG proteins interact, that MED12 and NANOG have similar DNA-binding profiles, and that MED12 promotes *Nanog* expression (Apostolou et al., 2013; Tutter et al., 2009). Several Mediator subunits, including *Med12*, have been identified in a screen for pluripotency maintenance that focused on transcription and chromatin regulators (Kagey et al., 2010). This study furthermore suggested that interactions between Mediator and cohesin contribute to genome folding and efficient enhancer-promoter interactions. When we probe *Med12* functions in lineage transitions, we find that, in contrast to *Med24,* loss of *Med12* does not compromise PrE differentiation (Hamilton et al., 2019). *Med12*-mutant cells however show a slower downregulation of pluripotency genes and a decreased ability to revert to naïve pluripotency in a colony-forming assay. We speculate that these cellular phenotypes are a reflection of a reduced ability of *Med12*-mutant cells to reconfigure the chromatin upon changing signaling environments. It is likely that the expression of individual genes is differentially sensitive to the loss of *Med12*. This may be the reason why expression from the *Sprouty4* locus, which is the most strongly upregulated gene upon acute FGF stimulation, shows a particularly high sensitivity to loss of *Med12*. In line with this idea, requirements of *Med12* for efficient induction are not exclusive to endogenous genes, but extend to exogenous transgenes such as the inducible GATA6-mCherry construct used to trigger primitive endoderm differentiation in our study. Surprisingly, such differential quantitative defects in the regulation of single genes upon loss of *Med12* do not lead to strong defects in acquiring early differentiated fates, such that transcriptomes of individual differentiated cells are not systematically different from each other in *Med12-*mutant and wild-type cells. This suggests that intracellular regulatory networks can buffer the composition of cellular transcriptomes against variable transcription efficiencies.

## Acknowledgements

We thank Kristin Nowak for generating *Fgf4*-mutant E14tg2a mESC lines, members of the Imig lab for help with lentivirus production, and the protein chemistry facility of the MPI Dortmund for supply of LIF and enzymes for molecular biology. Michelle Marten provided technical and organizational support for tissue culture and flow sorting throughout the project. We thank members of the Schröter lab, the Department of Systemic Cell Biology, Jochen Imig and Cristina Pina for critical discussions and feedback on earlier versions of the manuscript.

## Materials and Methods

### Cell culture

Routine culture of mESCs was performed at 37 °C with 5% CO_2_ in either serum + LIF medium (ESL, composed of GMEM with 10% fetal bovine serum (FBS), 2 mM GlutaMAX, 1 mM sodium pyruvate, 0.1 mM b-mercaptoethanol and 10 ng/mL LIF), on 0.1% gelatine coated dishes, or in 2i + LIF medium on fibronectin coated dishes. 2i + LIF is N2B27 supplemented with 1 µM PD0325901 (SelleckChem), 3 μM CHIR99201 (Tocris) and 10 ng/ml LIF (protein chemistry facility, MPI Dortmund). N2B27 was prepared as a 1:1 mixture of DMEM/F12 and Neuropan Basal Medium (both from PAN Biotech), supplemented with 1X N2 and 1X B27 supplements, 1X L-Glutamax, 0.0025% BSA, and 0.2 mM ß-mercaptoethanol (all from ThermoFisher). *Fgf4*-mutant cell lines were cultured in 2i + LIF supplemented with 10% FBS. Cells were passaged every two to three days, and detached with trypsin (PAN Biotech) or Accutase (Sigma-Aldrich).

### Cell lines

All cell lines generated in this study were derived from the E14tg2a wild-type line (Hooper et al., 1987). The GATA4-mCherry inducible line used for single cell RNA sequencing has been described previously (Raina et al., 2021). The *Spry4^H2B-Venus/+^*-reporter line was generated with a previously described targeting construct (Morgani et al., 2018) using lipofectamine 2000 according manufacturer’s instructions (Thermo Fisher Scientific). Correctly targeted clones were identified via long-range PCR as described in (Morgani et al., 2018). GATA6-mCherry inducible lines were established as described for GATA4-mCherry inducible lines in (Raina et al., 2021), but replacing the *Gata4* with a *Gata6* coding sequence in the PiggyBac vector for inducible gene expression. We established multiple clonal lines and tested them for GATA6-mCherry induction levels upon dox-treatment by flow cytometry. Three independent clones with induction levels similar or slightly higher than the previously established GATA4-mCherry inducible lines were selected for the experiment shown in Fig. 5D, and a single clonal line was chosen for all other experiments, including mutagenesis of *Med12*. Newly generated *Spry4^H2B-Venus/+^*-reporter and GATA6-mCherry inducible cell lines were checked for karyotypic abnormalities. To label nuclei for time lapse imaging cells were transfected with pCX-H2B-Cerulean-IRES-puro (Schumacher et al., 2024). Cell lines carrying PiggyBac transgenes were kept under appropriate selection to prevent transgene repression over passaging.

### sgRNA cloning and generation of single-gene mutants

For mutagenesis of individual genes via CRISPR/Cas9, gene targeting sgRNAs (Supp Table 7) were cloned into pX459 (Addgene plasmid #48139) using BbsI (NEB) overhangs following Ran et al., 2013 (Ran et al., 2013). Clonal mutant lines were generated using a combination of sgRNAs with targeting sequences 100 to 200 bp apart in the genome. Single sgRNAs were used when generating polyclonal lines. For validation experiments of the CRISPR screen (Figs. 1 and 2), we selected the most enriched sgRNA in sorted cells. A total of 1 µg of sgRNA containing px459 vectors was mixed with a final concentration of 0.04 µg/ml Lipofectamine 2000 (Thermo Fisher Scientific) in Opti-MEM (Gibco) according to the manufacturer’s protocol. For the generation of clonal lines, cells were seeded at clonal density into 10 cm dishes after transfection, for polyclonal experiments approximately 50k cells/cm^2^ were seeded. To enrich for successfully transfected cells, selection with 1.5 μg/ml puromycin was started 24 h after transfection for 48 h. To establish clonal lines, single-cell derived colonies were picked 4 to 6 d after transfection and expanded. For molecular characterization of genetic lesions, genomic DNA was purified with Terra™ PCR Direct Genotyping Kit (Takara), followed by PCR amplification and Sanger sequencing of specific genomic regions encompassing the target site.

### Genome-wide CRISPR Screen

To generate stably CAS9-expressing *Spry4^H2B-Venus/+^* reporter cells, cells were transduced with lentiCas9-Blast lentiviral particles (Addgene #52962-LV) at a multiplicity of infection of approximately 0.1. Transduction was performed with attached cells, 20 h after seeding at 20 000 cells/cm^2^, in presence of 5 µg/ml Polybrene in ESL. Continuous blasticidin (15 µg/ml, Gibco) selection was started 24 h after transduction. Lentiviral particles of the genome-wide gRNA library Brie (Addgene #73633) were generated according to standard protocols (Doench et al., 2016). For library transduction, 150 * 10^6^ CAS9-expressing *Spry4^H2B-Venus/+^* reporter cells were detached and mixed with the virus library in ESL with 5 µg/ml Polybrene. The following day, the same number of cells was reseeded and put under selection with puromycin (1.5 µg/ml, Sigma-Aldrich). Comparing cell counts with and without selection indicated a transduction efficiency of 25%, resulting in a >400-fold coverage of transduced cells per gRNA. In all subsequent steps, at least 31 * 10^6^ cells were processed to maintain gRNA coverage. To identify gRNAs enriched in cell populations with high and low *Spry*4:H2B-Venus expression, at least 0.5 * 10^6^ cells with the lowest or highest 1% of *Spry*4:H2B-Venus fluorescence, or 3 * 10^6^ cells with the lowest or highest 5% of *Spry*4:H2B-Venus fluorescence were FAC sorted and their DNA isolated by column-based genomic DNA purification (Monarch Genomic DNA Purification Kit, NEB). For reference, the genomic DNA of 31 * 10^6^ non-sorted control cells was purified in parallel. The integrated gRNA was PCR amplified using Pfu polymerase (prepared in house) with a sample specific, sequencing adapter and index containing primers (Supp Table 7; Carlini et al., 2021) using the complete purified genomic DNA as template. PCR-samples were purified with the SPRIselect reagent (Beckman Coulter) with double-sided size selection. Briefly, 0.5x SPRIselect was added to each sample, incubated for 5 min at RT and the SPRIselect removed with a magnet. This supernatant was again mixed with 1.2x SPRIselect, incubated and then discarded. After washing the beads, the DNA library was eluted from the beads and used for sequencing.

Paired-End Illumina Sequencing with a read length of 150 bp pairs was performed with at least 10 * 10^6^ reads per sorted sample and 30 * 10^6^ reads for the unsorted library controls. The raw reads were trimmed using Cutadapt (Martin, 2011) to remove the vector binding sequence. The reads were mapped to individual gRNAs, counted using *norm-method total* and statistically tested on the targeted gene levels using *gene-lfc-method alphamean* with Mageck (W. Li et al., 2014). Hits were selected based on the false discovery rate.

### Immunostaining

Immunostaining was performed as previously described (Schröter et al., 2015), Briefly, cells were washed with PBS containing Calcium and Magnesium, followed by fixation with 4% paraformaldehyde (Histofix, Sigma-Aldrich) for 15 min. Cells were permeabilized and blocked by rinsing and washing three times with PBS with 0.1% Triton X-100 and 1.0% bovine serum albumin (PBT-BSA). Primary antibodies (Anti-mouse NANOG (Affymetrix eBioscience, Cat.:14-5761), Anti-SOX17 (R&D systems, Cat.: AF1924), anti-Oct3/4 (POU5F1, Santa Cruz Biotechnology, Cat:. sc-5279)) were diluted 1:200 in PBT-BSA and incubated with the cells overnight at 4 °C. The next day, cells were washed in PBT-BSA and incubated with Alexa Fluor-conjugated secondary antibodies at 4 µg/ml (Invitrogen/Life Technologies) and Hoechst 33342 at 1 µg/ml (Invitrogen) in PBT-BSA in the dark for 2 h. Finally, samples were rinsed and washed with PBS and imaged in a mounting medium consisting of 80% glycerol, 16% PBS and 4% n-propyl-gallate.

### Immunoblotting

For western blot analysis of MED12 and ppERK, cells were washed twice with ice-cold PBS, supplemented with 1 mM activated orthovanadate in case of ppERK detection. Cells were mechanically detached in lysis buffer, based on commercially available lysis buffer (Cell Signaling) supplemented with benzonase (Sigma-Aldrich), cOmplete EDTA-free protease inhibitor cocktail (Roche), phosphate inhibitors P1 and P2 (Sigma). The lysates were snap frozen in liquid nitrogen twice and centrifuged. Protein concentration in the supernatant was measured with a micro BCA assay (Thermo Scientific). For western analysis, 20 µg of protein per sample were denatured by adding 5x Laemelli buffer and incubation at 95 °C for 5 min. The SDS-PAGE was run in 1x MOPS buffer (ThermoFisher) with 5 mM sodium-bisulfate and immediately transferred onto methanol-activated PVDF membranes. Transfer was performed in transfer buffer (12mM Tris-Base, 96mM Glycine, 20% methanol) at 40 V for 1.5 h in a NuPage transfer system (ThermoFisher). Membranes were blocked at RT for 1 h in Intercept blocking buffer (LI-COR), which was also used for the dilution and incubation with the primary antibodies anti-Tubulin 1:10000 (T6074, Sigma), anti-pERK1/2 1:1000 (4370S, Cell Signaling), anti-total ERK1/2 1:1000 (ab36991, Abcam) or anti-Med12 (1:250, A300-774A, Bethyl Laboratories). Appropriate secondary antibodies (IRdyes, LI-COR) were used at a dilution of 1:5000. Blots were imaged on an Odyssey CLx (LI-COR). Quantification of ppERK bands was performed using the gel quantification tool in Fiji, using the combined intensities of ERK1 and ERK2 bands, and normalizing ppERK by total ERK signals.

### In situ HCR

In situ hybridization probes against Spry4 and Nanog were designed by, and all commercial reagents for the staining procedure were obtained from Molecular Instruments, if not indicated otherwise. mRNAs were stained in solution as previously reported (Choi et al., 2018). Specifically, cells from confluent 6-well plate wells were washed with PBS and detached using trypsin. Following centrifugation for 5 min at 200xg, the specification for all further centrifugation steps, cells were fixed with 4% PFA for 1 h. After removal of the fixative via centrifugation, cells were washed four times with PBS with 0.1% Tween20 (PBST, Bio-Rad). Cells were incubated overnight in ice-cold ethanol. Starting with 1 * 10^6^ cells, and two additional washes with PBST, cells were incubated in 400 µl probe hybridization buffer for 30 min at 37 °C. 2 pmol of gene-specific hairpins were prepared in 100 µl preheated probe hybridization buffer and added to the cell suspension, which was subsequently incubated overnight at 37 °C. Preheated wash buffer was used to remove residual hybridization oligos by washing thrice with 10 min of intermediate incubation. One final wash and 5 min incubation were performed in saline-sodium citrate (SSC, Sigma-Aldrich) buffer with 0.1% Tween20. Cells were resuspended in 150 µl amplification buffer. 10 pmol of each labeled hairpin were mixed, heated to 95 °C for 90 s, and cooled back to RT. Hairpins were added into 100 µl of amplification buffer, mixed with the cell suspension and incubated in the dark for 1 h. A final five washes with SCCT were followed by mounting the sample onto a microscopy glass slide. Samples were resuspended in 20 to 100 µl SCCT to ensure high single-cell densities. 2 µl of the cell suspension were squished between the slide and a cover slip to maximize area per cell and distance between mRNA spots.

### RNA labeling

Newly synthesized RNA was labelled using the Click-iT™ RNA Alexa Fluor™ 594 Imaging Kit (Invitrogen) according to the manufactures protocol. Briefly, we incubated cells with 1 mM 5-ethynyl uridine for the last 30 min or 1h of differentiation or during pluripotency. After, washing, fixation with 4% PFA and permeabilization, the click reaction with the Alexa Fluor 594 azide was performed for 30 min, followed by washing and nuclear staining with SiR-DNA 647 (Spirochrome) for 15 min.

### Imaging and Image Analysis

Tilescans of immunostainings were imaged with a Leica SP8 confocal microscope (Leica Microsystems) with a 63x 1.4 NA oil immersion objective. Images were analyzed in Fiji (Schindelin et al., 2012). For segmentation, StarDist 2D (Schmidt et al., 2018) was used using the versatile (fluorescent nuclei) model and default post processing parameters. Mean fluorescence intensity was measured in segmented cells in all acquired channels. Cells with a nuclear area smaller than 40 µm^2^ were filtered out. To determine fluorescence intensity threshold values for the classification of cell types, we manually selected thresholds that best bisected the bimodal expression profiles of the lineage markers. The same thresholds were applied to different samples in a single experiment. In the RNA-labeling experiment, after segmentation based on the nuclear staining and intensity measurement, a gaussian mixture model was fitted to the intensity distributions of the H2B-Cerulean channel. Using the posterior probability, the fluorescence intensity thresholds at which a cell can be assigned to the correct genotype with an ≥ 85% certainty were determined and genotype assignment was performed accordingly, not assigning cells in between the populations. The integrated intensity of the RNA-AF594 signal within the nucleus region was used to determine the amount of newly produced RNA within the labeling period.

Images of live *Spry4^H2B-Venus/+^*-reporter cells were acquired with an Olympus IX81 widefield microscope, equipped with a stage top incubator (ibidi), pE4000 illumination (CoolLED), ORCA-Quest qCMOS camera (Hamamatsu) with a 63x 1.35 NA oil immersion objective. Hardware was controlled by Olympus CellSens Software. Images of in situ HCR for mRNA counting were acquired on the same system with a 100x 1.4 NA objective with 1024×1024 pixels and at least 20 z-slices, separated by 0.4 µm. Single cells were manually segmented on a brightfield image, taken in the middle of the z-stack. Counting mRNA spots from in situ HCR was performed using RS-FISH (Bahry et al., 2022). The anisotropy was determined from single spots and set to 1.2. No robust fitting was applied. Sigma was set to 0.994. The threshold, the image maximum and the minimal spot intensity threshold were set for each channel per replicate manually, to account for different (background) intensity levels. Time lapse imaging was performed with a 40x 0.9 NA objective on an Olympus IX81 widefield microscope, equipped with an LED-based illumination system (pE4000, CoolLED) and an iXon 888 EM-CCD camera (Andor). MicroManager (Edelstein et al., 2010) was used to control the hardware. Images were taken every 10 min. Tracking was performed with the manual tracking function in Trackmate v7 (Ershov et al., 2022) and fluorescence intensity was measured as the mean intensity in a spot with a 4 µm radius within the nucleus. In R, tracks were smoothed with a rolling average over 7 frames. For ROC analysis the R package pROC (Robin et al., 2011) was applied and the optimal threshold was defined by the Youden’s J statistic (Youden, 1950).

### Flow cytometry

Analysis of *Spry4*:H2B-Venus reporter expression in live or fixed cells was performed on a LSRII flow cytometer (BD Biosciences). Cell sorting and analysis of GATA6-mCherry expression was carried out using a FACS Aria Fusion (BD Biosciences). Primary data analysis including gating single cells based on SSC and FSC was done with FlowJo version 9 (BD Biosciences).

### Clonogenicity Assay

Clonogenicity assays were performed according to (Kalkan et al., 2017). Briefly, 1 * 10^4^ cells/cm^2^ were seeded in 2i + LIF for 24 h, followed by differentiation in N2B27 for 48 h. Control wells for each parental cell line were kept in 2i + LIF. Cells were then detached with Accutase to single cells, and 500 cells were reseeded into 6-well plates with 2i + LIF + 10% FBS. 10% FBS were included to support survival of *Fgf4*-mutant cells.

After 5 d, the colonies formed were fixed and stained with an alkaline phosphatase assay kit (Sigma-Aldrich) to distinguish pluripotent and differentiated colonies. Tile scans of the wells were acquired with an Olympus IX81 widefield microscope with a 4x 0.16 NA objective. We applied background subtraction, gaussian blurring, Otsu-thresholding, and conversion of images to a binary mask in ImageJ, and then used the *AnalyzeParticles* function to set thresholds for size and circularity and to determine the number of colonies. Colony numbers were normalized to the number of colonies obtained in the control.

### Bulk RNA sequencing

For bulk RNA sequencing, cells were seeded at a density between 3.5 and 5.5 * 10^4^ cells/cm^2^ in 2i or N2B27 + Chiron + LIF, followed by stimulation under indicated conditions. Replicates were obtained either from independent biological experiments (Fig. 3 A, B) or from both independent biological experiments and independent *Med12-*mutant lines (Fig. 3F, G). RNA isolation was performed with TRIzol (ambion) according to the manufacturer’s instructions. Sequencing libraries were prepared on polyA-enriched RNAs, followed by paired-end sequencing at a read-length of 150 bp and depth of approximately 30 * 10^6^ reads per sample. Strand-specific libraries were generated only for the FGF-titration experiment (Fig. 1 Supp 1) and the differentiation of the *Med12* wild-type and mutant cells (Fig. 3A, B). Raw reads were mapped to the mouse genome (GRCm39, release 108 (both *Med12* mutant experiments) or release 97 (FGF titration experiment) with hisat2 (v2.1.0; Kim et al., 2019). SeqMonk was used to quantify counts per gene, either as TPM or as raw counts as input for downstream DESeq2 analysis (Love et al., 2014) for identification of differentially expressed genes.

Differentiation delay in Med12 mutants was estimated according to (Lackner et al., 2021). We first determined the expression change of the naïve marker genes *Nanog*, *Esrrb*, *Tbx3*, *Tfcp2l1*, *Klf4*, *Prdm14* and *Zfp4* in *Med12*-mutant and wild-type cells, and then plotted the Euclidean distance of this expression change to that of the time-resolved dataset from Lackner et al., 2021.

Signaling footprint analysis in *Med12* mutants was performed similarly to Lacker et al., 2021. This study defined a specific set of target genes for each pluripotency associated signaling system based on gene expression changes in knockouts of signaling genes. A signaling footprint for a knockout line can then be determined from the difference in the expression of pathway footprint genes to the wild-type line after 24 h of differentiation. Measures for the signaling footprint are the Spearman correlation between each knockout line and the respective pathway defining knockout, and the ratio between the sum of expression fold changes between a knockout line and the respective pathway defining knockout, defined as pathway activity. To compare the *Med12* mutant data from this study, the wild type conditions were used for batch correction.

### Cell multiplexing and scRNA sequencing

Cells for scRNAseq were seeded at a density of 3.5 * 10^4^ cells/cm^2^ in 6-well plates in 2i + LIF and grown overnight. The next morning, induction in 2i + LIF + dox was first started in the mutant clones, and 4 h later in the wild-type lines. After 8h and 4 h, respectively, induction was stopped by washing once with N2B27, followed by 20 h of differentiation in N2B27. Controls for each cell line were continuously kept in 2i + LIF. For sequencing, cells were washed three times with PBS and detached with Accutase. Accutase was removed by centrifugation and 1 * 10^6^ cells per sample were resuspended in PBS + 0.04 % BSA and immediately used for multiplexing labeling following the protocol of 10x Genomics for samples with a viability above 80 % (Cell Multiplexing Oligo Labeling for Single Cell RNA Sequencing Protocols with Feature Barcode technology, CG000391). Briefly, cells were spun down once again, resuspended with individual cell multiplexing oligos (CMO no. 301 to 310) and incubated for 5 min at RT. Cells were washed twice with PBS + 1 % BSA and passed through a cell strainer (FALCON, mesh size 35 µm). A total of 1.2 * 10^5^ single cells from all samples were pooled at equal ratios, and 4 * 10^4^ were used for droplet generation, corresponding to a target number of 2.4 * 10^4^ cell-containing droplets. Droplet generation, lysis, mRNA and cell barcode capture, and generation of both the gene expression library as well as the cell multiplexing library was performed following the instructions by 10x genomics (Chromium Next GEM Single Cell 3ʹ Reagent Kits v3.1 (Dual Index) with Feature Barcode technology for Cell Multiplexing, CG000388). Specifically, we chose 11 PCR cycles for cDNA amplification and 10 cycles for the sample index PCR. Concentration and insert size distribution for both the gene expression library and the cell multiplexing library were determined with a BioAnalyzer High Sensitivity DNA Assay (Agilent). Sequencing was performed on a NovaSeq 6000 on multiple flowcells with a paired-end 150 bp configuration. In total 1.2 * 10^9^ and 2.3 * 10^8^ read pairs were obtained for the gene expression and multiplexing library, respectively.

Demultiplexing to the individual samples, based on the cell multiplexing barcode and alignment to the mouse genome mm10 (GENCODE vM23/Ensembl 98, obtained from 10x Genomics) was performed with CellRanger (version 7.1.0, 10x Genomics). Downstream analysis was performed in R with Seurat v5 (Hao et al., 2023). We first filtered cells by removing barcodes with ≤ 2500 detected genes and ≥15 % of reads aligned to mitochondrial genes, retaining between 1100 and 1700 cells per sample with median mRNA counts per cell between 23233 and 27890 in the different samples. mRNA counts for each gene were normalized by dividing its counts by the total number of counts per cell, multiplied by 10000. Log1p transformation was applied before plotting expression data as violin plots. For downstream analysis and representation of gene expression as heatmaps, centering counts for each feature and scaling to its standard deviation was applied. Principal component analysis was performed on the 2000 most variable features in the relevant subset of cells. The resolution of the Louvain clustering algorithm was set to 0.05 when clustering multiple samples. In case of Jaccard-Index estimation the clustering resolution was set to 0.15 and the clustering was performed on each sample separately. 100-fold repetition of this clustering approach with a random subset of the data with 70 % of the cells allowed the calculation of a Jaccard-index, as previously described (Tang et al., 2021). For annotation of the Epi- and PrE-fate, the cells of the differentiated samples were integrated with Seurat integration based on the *rpca* reduction. Differentially expressed genes between cell states and genotypes were identified with the *FindMarkers* function in Seurat with a minimal expression difference in the log1p transformed expression values of 0.5. Biological noise was quantitated and distinguished from modeled technical noise in local neighborhoods of each cell with VarID2 (Rosales-Alvarez et al., 2023).

### Data and code availability

Sequencing data from this paper has been deposited at GEO with accession number GSE253609. All code used for analysis and visualization, together with a list of the R packages used, is available from the authors upon request.

## Supplementary Figures

**Fig. 1 Supp 1:**
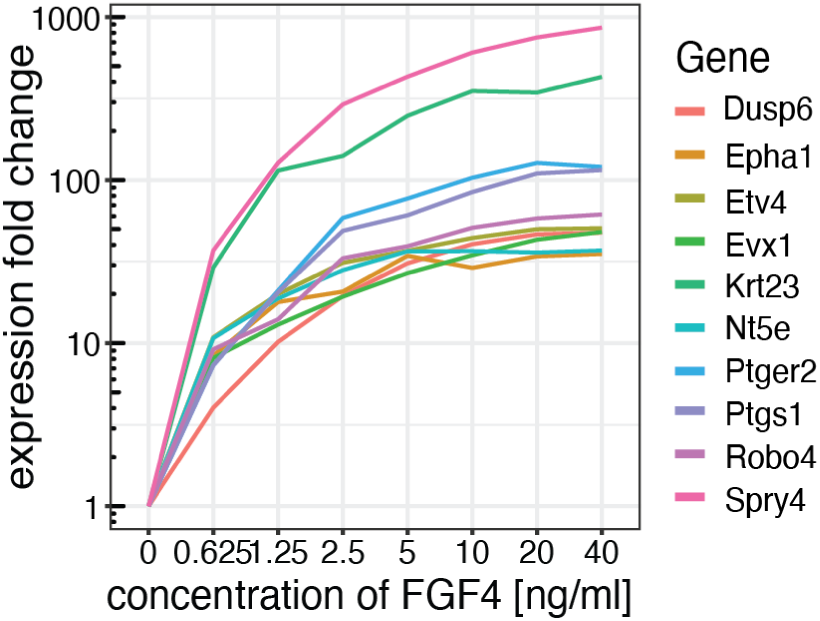
*Spry4* is strongly upregulated upon FGF stimulation. Expression fold change of the ten most upregulated genes upon FGF4 titration in *Fgf4* mutants. *Fgf4*-mutant *Spry4^H2B-Venus/+^* cells were transitioned from 2i + LIF medium containing 10% FBS to N2B27 supplemented with Chiron and LIF for 18 h, followed by 6 h of stimulation with indicated concentrations of FGF4 in N2B27 with Chiron.

**Fig. 1 Supp 2:**
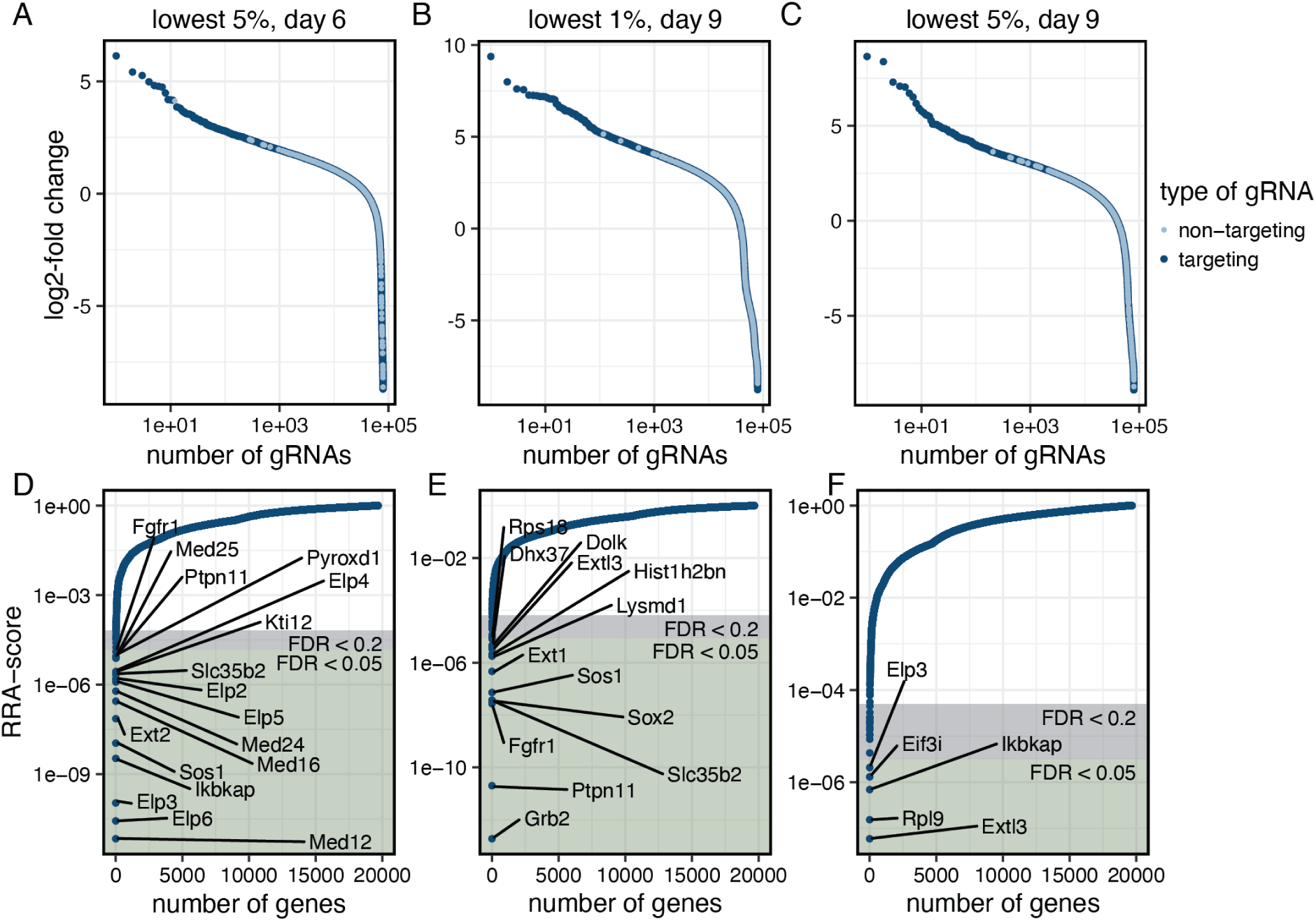
Genome-wide CRISPR knockout screen results in robust enrichment of gRNAs and corresponding genes. **A - C** Log 2-fold enrichment of gene-targeting (dark blue) and control gRNAs (light blue) in cells sorted for the lowermost 5% of H2B-Venus signal on day 6 after gRNA transduction (A), sorted for the lowermost 1% on day 9 after transduction (B), or sorted for the the lowermost 5% on day 9 after transduction (C). **D** - **F** RRA scores for genes corresponding to enriched gRNAs identified in A - C.

**Fig. 2 Supp 1:**
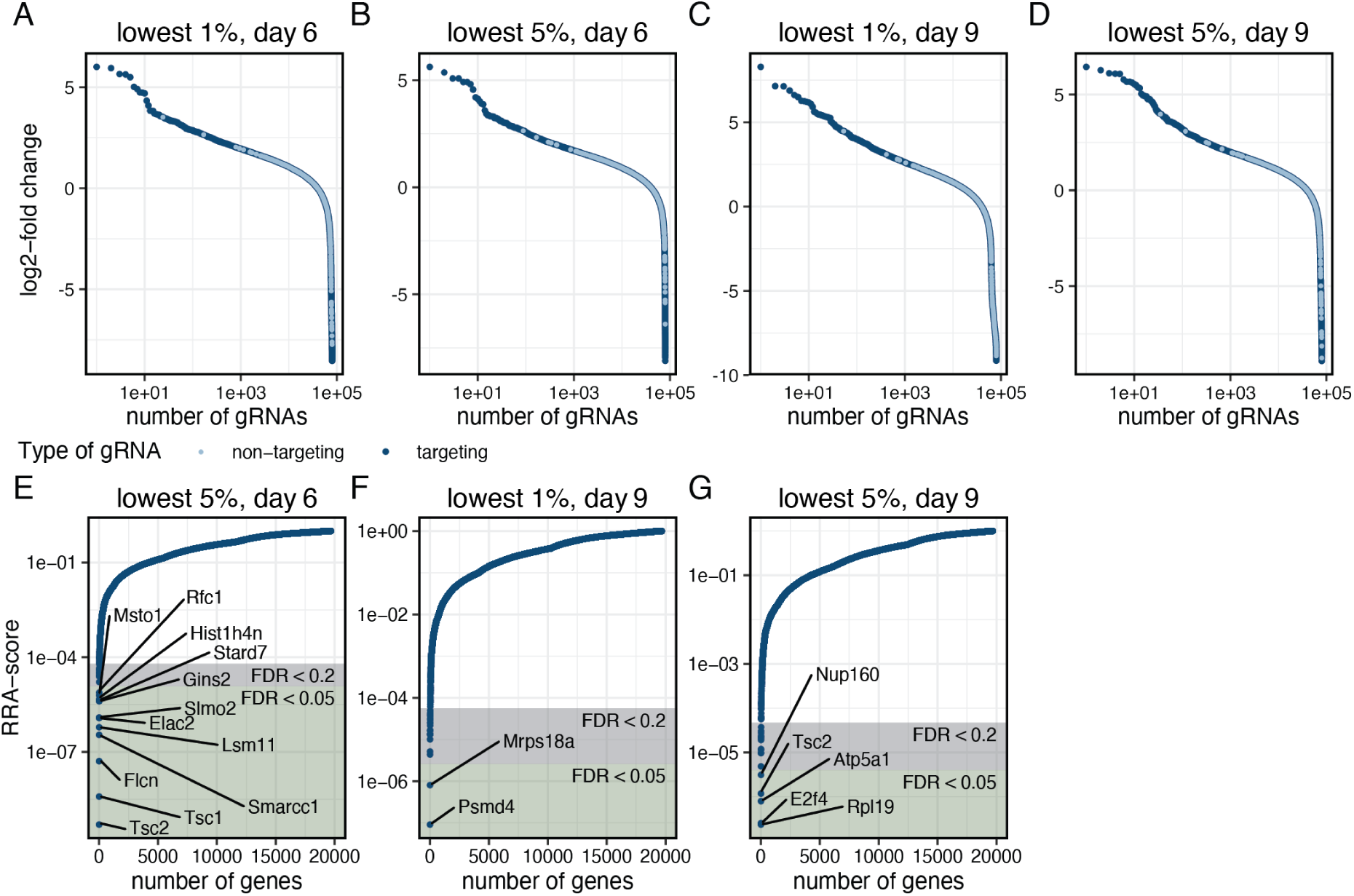
Robust enrichment of gRNAs and corresponding genes that negatively regulate Spry4:H2B-Venus expression. **A** - **D** Log 2-fold enrichment of gene-targeting (dark blue) and control gRNAs (light blue) in cells sorted for high H2B-Venus expression on different days as indicated. **E** - **G** RRA scores for genes corresponding to enriched gRNAs identified in B - D. RRA scores for genes corresponding to gRNAs enriched in the 1% of the cells with highest fluorescence after 6 days are shown in Fig. 2A.

**Fig. 3 Supp1:**
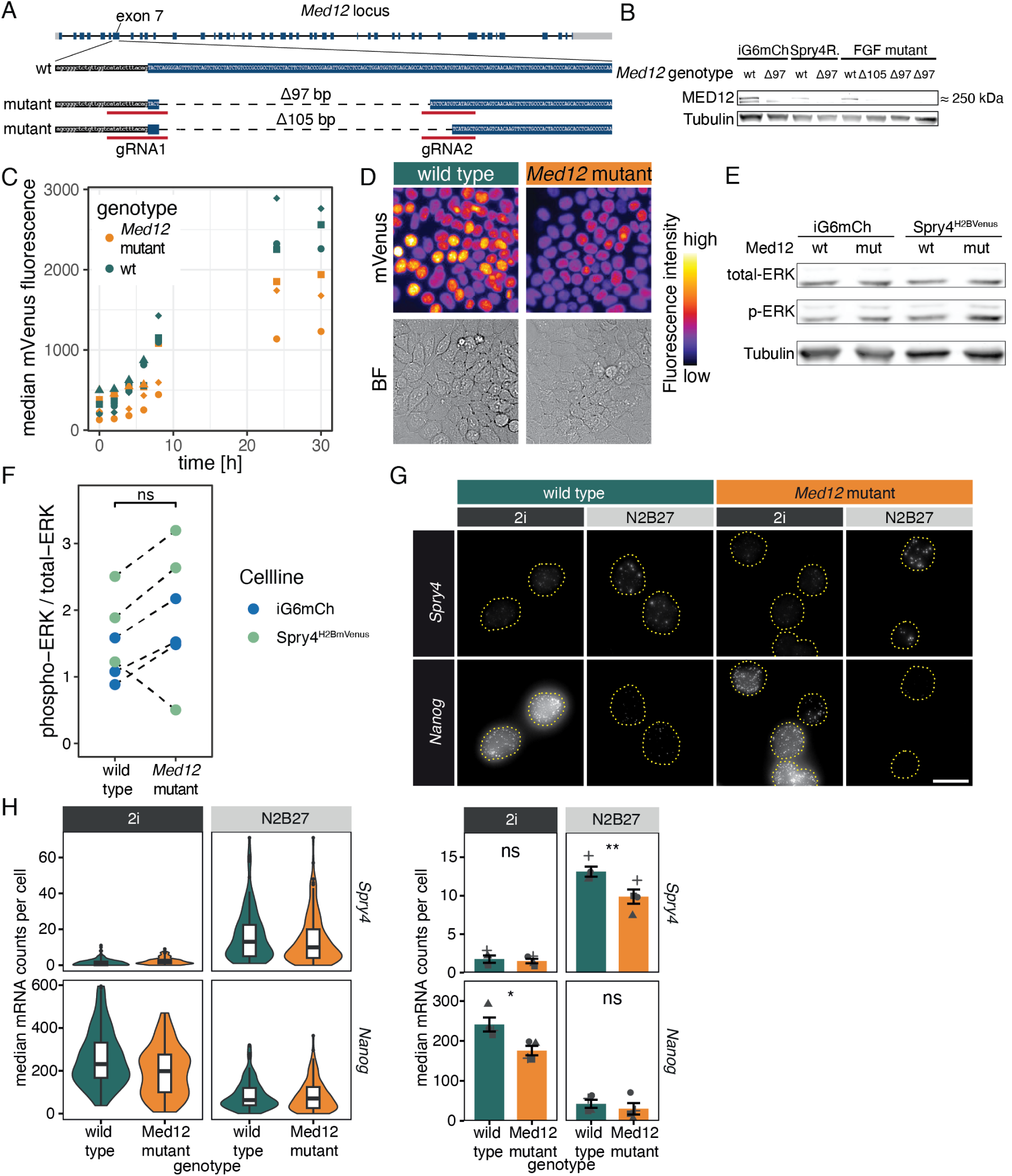
Generation of *Med12* mutant cell lines. **A** Schematic of the *Med12* gene locus and the gRNAs used to create a *Med12* loss-of-function by deleting 97 or 105 bp of exon7. Intronic sequence in lowercase and black underline, exonic sequence in uppercase and blue underline. **B** Immunoblotting for MED12 and Tubulin in cell lysates from independent monoclonal *Med12* mutant lines generated in different genetic genetic backgrounds. Δ97 and Δ105 indicate the type of exon7 deletion in each clone. **C** *Spry4^H2B-Venus/+^* expression upon release from 2i + LIF to N2B27 in wild-type and *Med12*-mutant cells measured by flow cytometry. Data points show median fluorescence in each experiment. N = 3. **D** H2B-Venus expression in live wild-type and *Med12-*mutant *Spry4^H2B-Venus/+^* cells after 24 h of growth in N2B27 following release from 2i + LIF. **E** Immunoblotting of cell lysates from *Med12* wild type and *Med12-*mutant *Spry4^H2B-Venus/+^* and iGata6 mESCs, stained for Tubulin, total- and phopsho-ERK. **F** Quantification of phospho-ERK signals from immunoblots, normalized to total-ERK. N=3. ns indicates p ≥ 0.05, paired two-sided t-test. **G** Fluorescent in-situ hybridization staining of wild-type (left) and *Med12*-mutant cells (right) for *Spry4* (top) and *Nanog* (bottom) mRNAs in 2i medium, or 24h after transfer into N2B27. **H** Violin plots (left) and summary statistics of *Spry4* (top) and *Nanog* (bottom) mRNA numbers in wild-type and *Med12* mutant cells determined by in-situ hybridization as described in **G.** Violin Plots show one out of N = 4 replicates, n > 100 cells per replicate and condition.

**Fig. 3 Supp 2:**
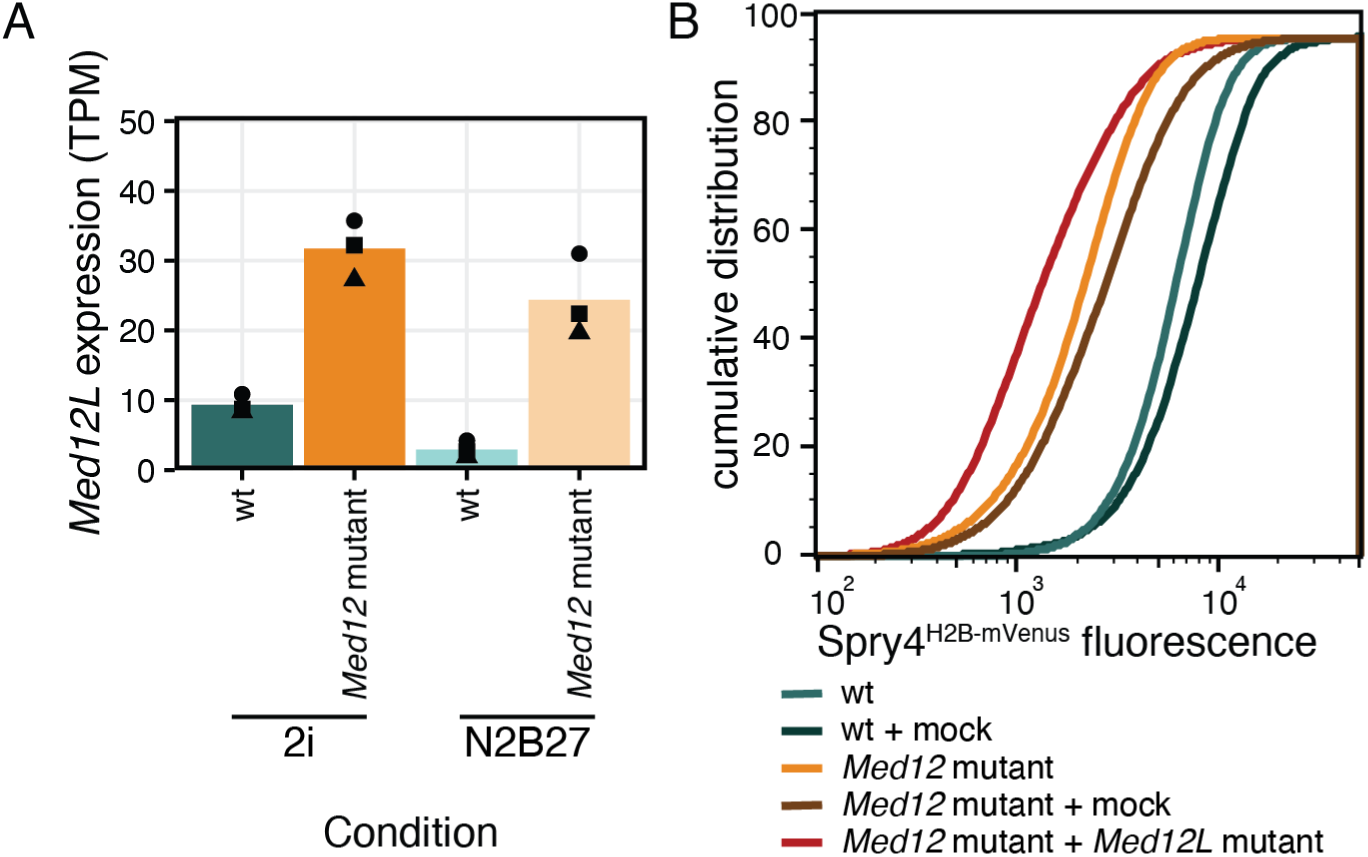
*Med12l* is upregulated upon loss of *Med12*. **A** Expression of *Med12l* in wild-type and *Med12*-mutant cells (data from RNA-seq experiment in Fig. 3A). **B** H2B-Venus expression in untransfected wild-type and *Med12*-mutant cells, and after transfection with mock or *Med12l-*targeting gRNAs. Expression was measured by flow cytometry 7 d after transfection and culture in ES + LIF medium.

**Fig. 4 Supp 1:**
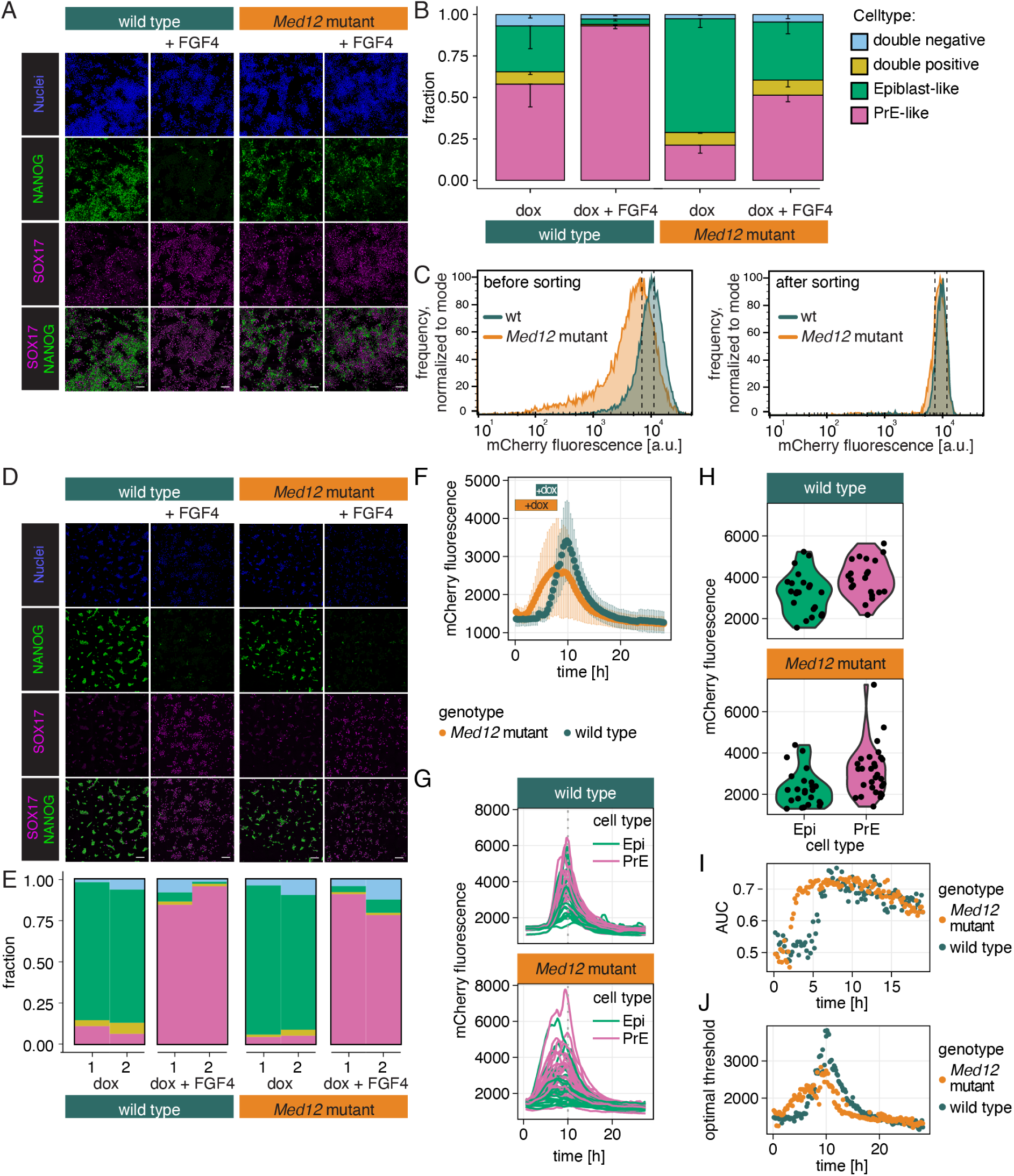
PrE differentiation in *Med12* mutant cells. **A** Immunostaining of the Epi-marker NANOG (green) and the PrE marker SOX17 (magenta) after 8 h of GATA6 induction and 20 h of differentiation with and without exogenous FGF4 in wild-type and *Med12*-mutant cells. Scale bar: 100 µm. **B** Cell type proportions in wild-type and *Med12*-mutant cells differentiated as in (A). N=3, n > 1100 cells per replicate, error bars indicate SEM. **C** Gata6-mCherry fluorescence after 8 h of dox induction measured by flow cytometry. Left shows distribution of expression levels in the whole population, right shows expression levels after flow sorting of cells with similar fluorescence intensity. Dashed lines indicate sorting gate. **D** Immunostaining of the Epi-marker NANOG (green) and the PrE marker SOX17 (magenta) after 8 h of GATA6 induction, flow sorting as described in (C), reseeding and 20 h of differentiation with and without exogenous FGF4 in wild-type and *Med12*-mutant cells. Scale bar: 100 µm. **E** Cell type proportions in wild-type and *Med12*-mutant cells differentiated as in (D). N=2, n > 500 cells per replicate. **F** Quantification of Gata6-mCherry expression dynamics from time-lapse movies during induction and differentiation. Boxes indicate induction times (8 h for *Med12* mutant, 4 h for wild type). Error bars indicate SD. One out of N = 5 replicates shown, n > 300 cells per time point. **G** Same experiment as in (E), but showing Gata6-mCherry fluorescence in single cells. Trace color indicates differentiation outcome determined by immunostaining (Epi: green; PrE: magenta). **H** Gata6-mCherry fluorescence in single cells 2 h after the end of induction, plotted separately for prospective Epi and PrE cells. **I** Predictive power of GATA6-mCherry expression determined as Area Under the Curve (AUC) from ROC-analysis. **J** Optimal GATA6-mCherry threshold to predict differentiation outcome determined by Youden’s J statistic of ROC-analysis. **F** - **I** Data from one representative experiment out of N = 3 replicates, n ≥ 45 cells per genotype.

**Fig. 4 Supp 2:**
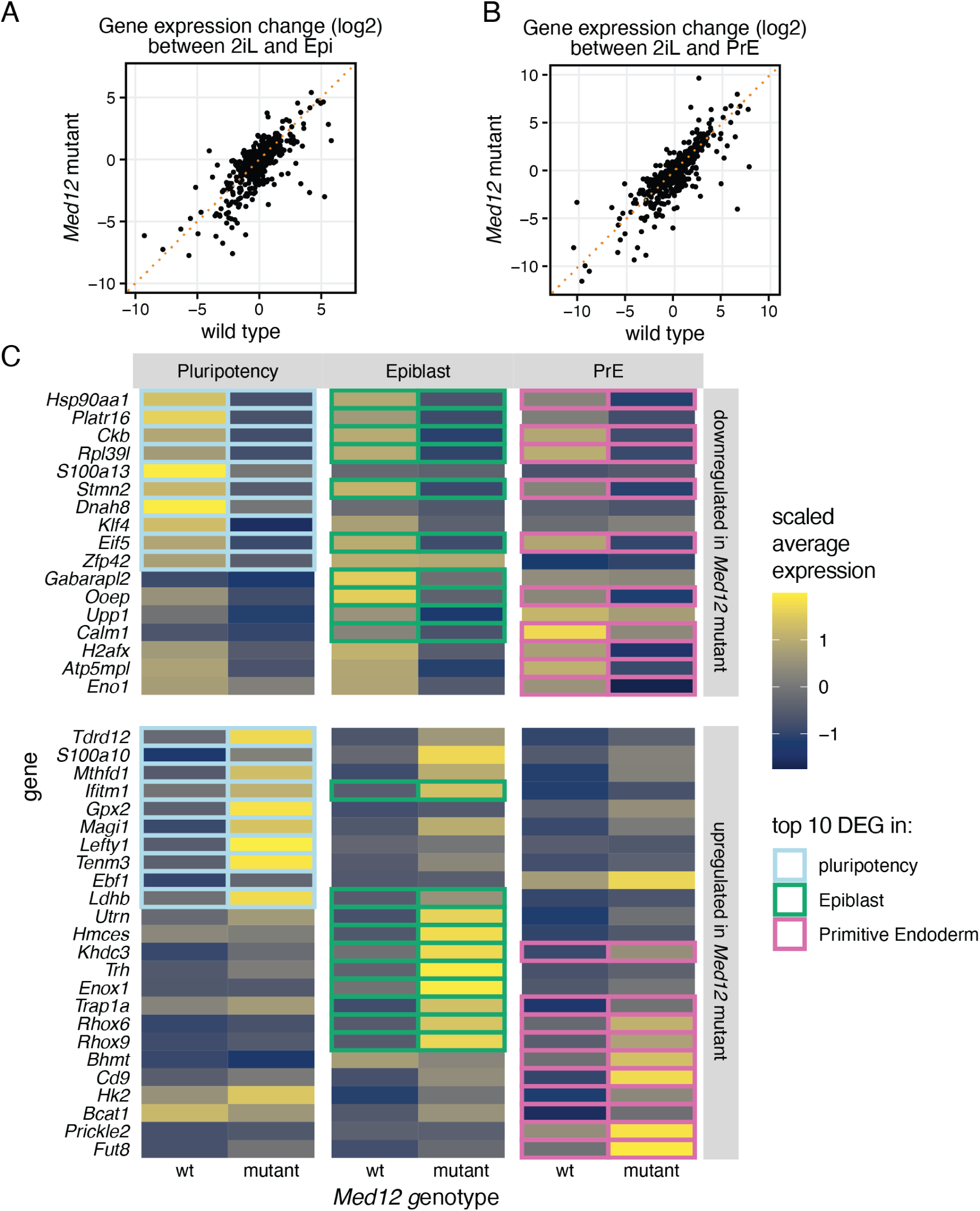
**A** and **B** Expression change of each gene upon differentiation from pluripotency (2iL) to Epi (A) and PrE (B) in wild-type versus *Med12*-mutant cells. Dotted, orange line indicates the unity line. **C** Differentially expressed genes between *Med12* wild type and mutant cells for the three different cell states. Tile color shows scaled average gene expression, colored boxes indicate the 10 genes with the largest fold-change between *Med12* wild type and mutant cells in each cell state.

**Fig. 4 Supp 3:**
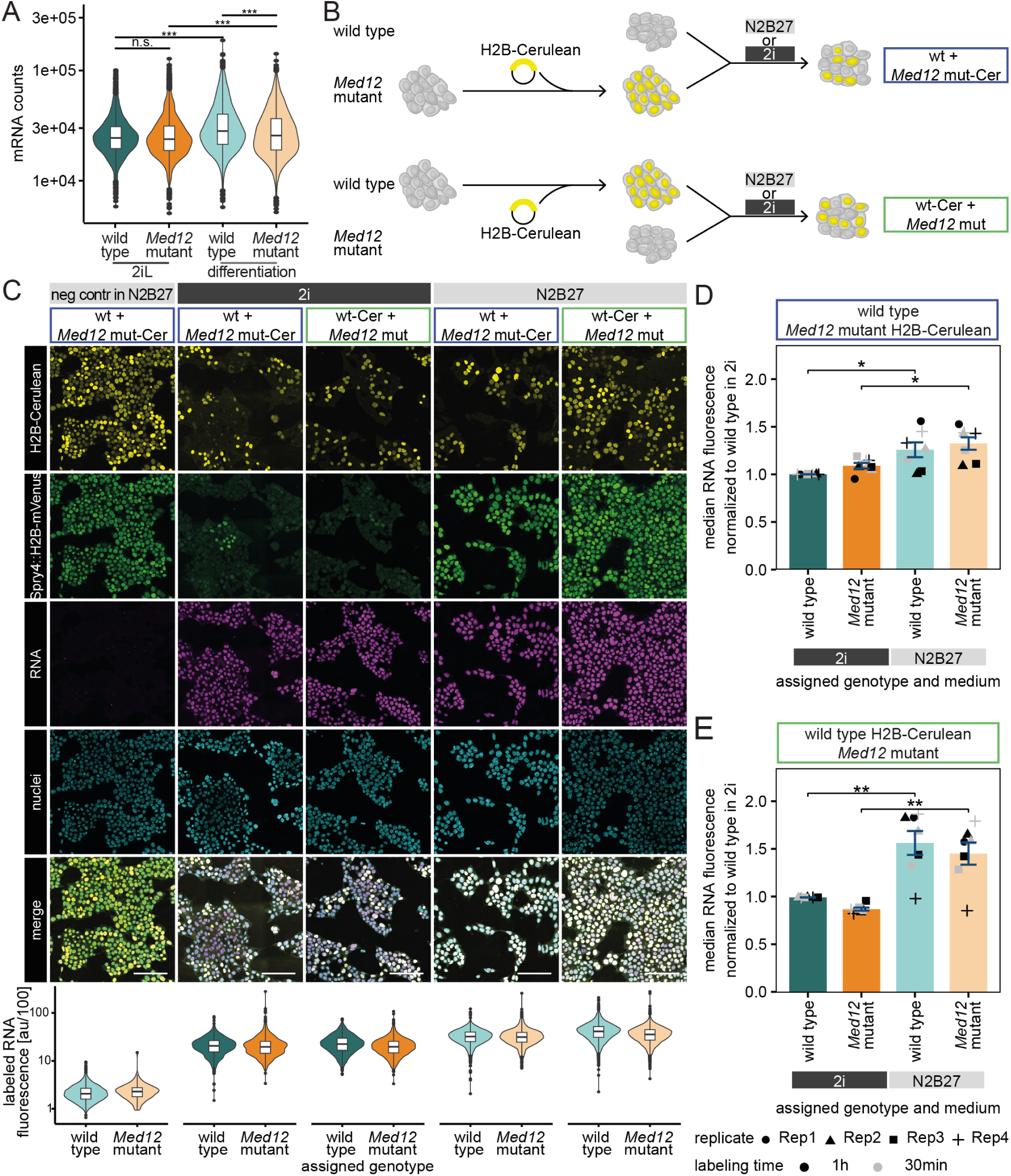
Global RNA levels increase during differentiation of mESCs. **A** Raw number of UMIs per cell detected by scRNAseq in wild-type and *Med12*-mutant cells. ns indicates p ≥ 0.05, *** indicates p ≤ 0.001, Bonferroni-adjusted Kolmogorov-Smirnov test. **B** Schematic of H2B-labeling of *Med12* mutant (top) or wild type cells (bottom) to compate RNA production in pluripotency (2i) and differentiation (N2B27) conditions. **C** Fluorescent images (upper panel) and violin plot of intensity (lower panel) of RNA-labeling with EU in 2i (dark grey) and N2B27 (light grey). Mixtures of genotypes were images in the same well, with one of them labeled with H2B-Cerulean (yellow) expression. For nuclei segmentation hoechst staining (cyan) was used and Spry4-H2B-mVenus-expression (green) confirmed differentiation in N2B27. **D, E** Quantification of intensity of RNA labeling based on cells assigned genotype based on their H2B-Cerulean expression (B). In (C) the *Med12*-mutant cell line was labeled with H2B-Cerulean, in (D) the wild-type line. N = 3, n > 750 cells. n.s. indicates p ≥ 0.05, * indicates p < 0.05 and ** indicates p < 0.01, paired two-sided t-test with Bonferroni multiple testing correction.

**Fig. 5 Supp 1:**
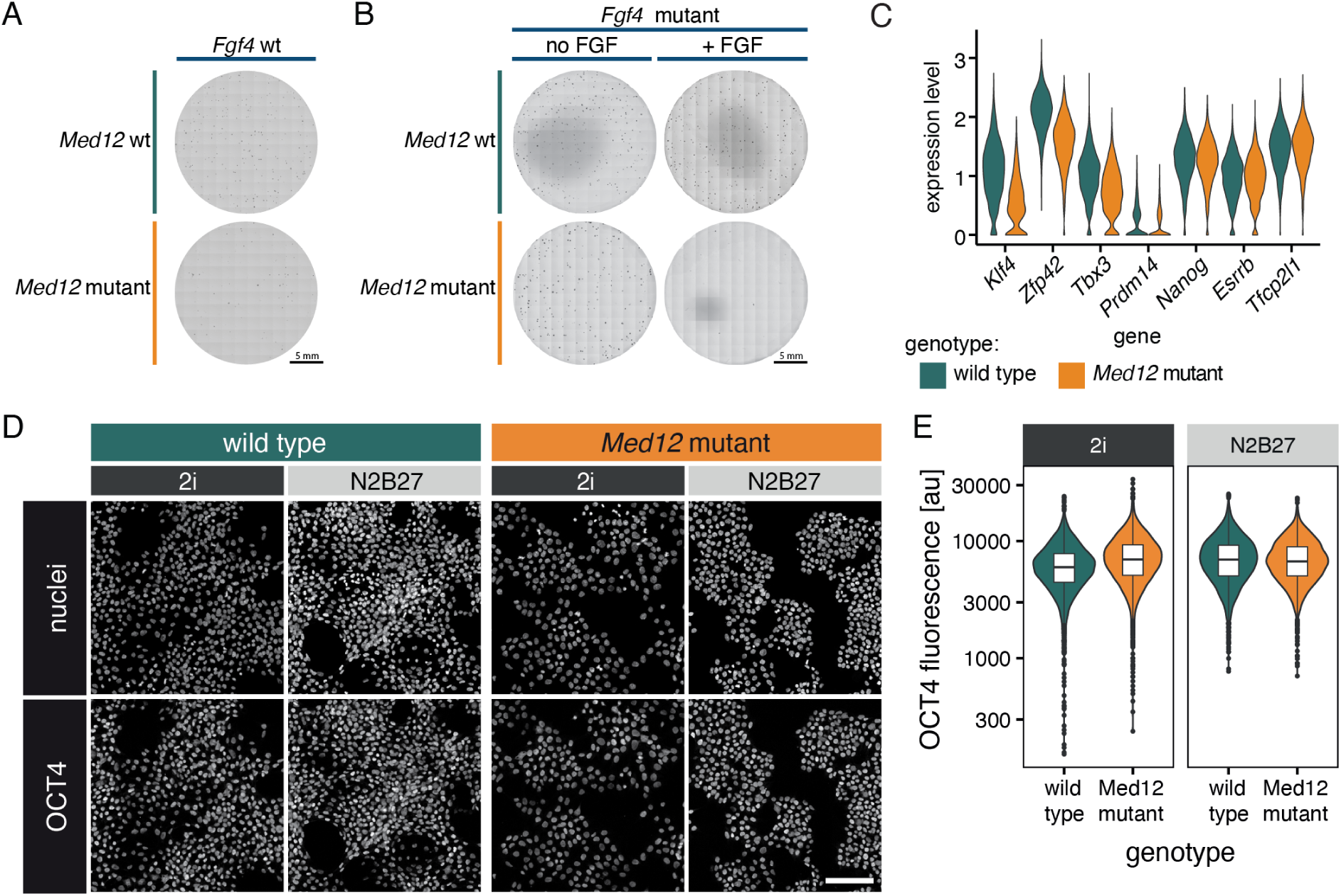
Reduced clonogenicity in *Med12* mutants compared to wild type. **A** and **B** Representative images of plates from the clonogenicity assay depicted in Fig. 5A, corresponding to quantifications shown in Fig. 5B. **A** shows plates with wild-type and *Med12-*mutant cells in an *Fgf4* wild type background, **B** shows plates with wild-type and *Med12-*mutant cells in an *Fgf4*-mutant background without (left) and with (right) FGF4 supplementation. **C** Ln-transformed expression levels of same naïve pluripotency marker genes as in Fig. 5C,D from pluripotent cells of single cell sequencing experiment from Fig 4. **D** Immunostaining for OCT4 (Pou5f1) in wild-type and *Med12*-mutant cells in 2i and N2B27. Scale bar 100 µm. **E** Quantification of OCT4 fluorescence per nucleus from cells stained as in (D).

## Supplementary Table legends

**Supp Table 1:** Raw and processed data from CRISPR screen including detected counts of gRNAs and enriched genes.

**Supp Table 2:** Differentially expressed genes comparing wild-type and *Med12*-mutant cells in 2i and after 24 h differentiation in N2B27.

**Supp Table 3:** Differentially expressed genes in *Fgf4*-mutant and *Fgf4 Med12* double mutant cells upon FGF4 stimulation in N2B27.

**Supp Table 4:** Differentially expressed genes between the Epi- (cluster 1) and PrE-cells (cluster 0) determined by single cell RNA sequencing experiment.

**Supp Table 5:** Differentially expressed genes between wild-type and *Med12*-mutant cells separately in pluripotency conditions, the Epi- and PrE-cluster.

**Supp Table 6:** Raw and normalized counts of colonies detected in the colony formation assay.

**Supp Table 7:** Oligos used as gRNAs or PCR primers.

## Supplementary Movie legends

**Supp Movie 1**: Timelapse imaging of H2B-Cerulean (blue) and iGata6-mCherry expression (red) in wild-type and Med12-mutant cells during Epi and PrE differentiation under same conditions as in Figure 3E. Following time lapse imaging, cells were immunostained for the Epi-marker NANOG (green) and the PrE marker SOX17 (magenta). Nuclei in immunostainings labelled with Hoechst dye (cyan). Scale bar: 50 µm.

